# Revisiting the Notion of Deleterious Sweeps

**DOI:** 10.1101/2020.11.16.385666

**Authors:** Parul Johri, Brian Charlesworth, Emma K. Howell, Michael Lynch, Jeffrey D. Jensen

**Affiliations:** School of Life Sciences, Arizona State University, Tempe, AZ 85287, United States; Institute of Evolutionary Biology, School of Biological Sciences, University of Edinburgh, EH9 3FL, United Kingdom; Center for Mechanisms of Evolution, The Biodesign Institute, Arizona State University, Tempe, AZ, United States

## Abstract

It has previously been shown that, conditional on its fixation, the time to fixation of a semi-dominant deleterious autosomal mutation in a randomly mating population is the same as that of an advantageous mutation. This result implies that deleterious mutations could generate selective sweep-like effects. Although their fixation probabilities greatly differ, the much larger input of deleterious relative to beneficial mutations suggests that this phenomenon could be important. We here examine how the fixation of mildly deleterious mutations affects levels and patterns of polymorphism at linked sites - both in the presence and absence of interference amongst deleterious mutations - and how this class of sites may contribute to divergence between-populations and species. We find that, while deleterious fixations are unlikely to represent a significant proportion of outliers in polymorphism-based genomic scans within populations, minor shifts in the frequencies of deleterious mutations can influence the proportions of private variants and the value of *F_ST_* after a recent population split. As sites subject to deleterious mutations are necessarily found in functional genomic regions, interpretations in terms of recurrent positive selection may require reconsideration.

## INTRODUCTION

Among the most important results in theoretical population genetics, now nearly a century old, are the fixation probabilities of new beneficial and deleterious mutations, which were obtained by Fisher (1922, 1930), Haldane (1927) and Wright (1931), using different approaches. Their results were later generalized by Kimura (1957, 1962, 1964), using the backward diffusion equation. A somewhat lesser-known result concerns the trajectories of these selected mutations. Specifically, Maruyama & Kimura (1974) found that, conditional on fixation, the time that a beneficial autosomal mutation with selection coefficient +*s* and dominance coefficient *h* spends in a given interval of allele frequency in a randomly mating population is the same as that for a deleterious mutation with selection coefficient *−s* and dominance coefficient 1 − *h*, provided that the conditions for the validity of the diffusion equation approximation hold (*i.e*., the change in allele frequency of the mutation is small enough to be approximated by a continuous-time diffusion process). Thus, given that the effects of selective sweeps on variability at linked neutral sites are related to their speed of transit through a population (Maynard Smith & Haigh 1974; Stephan 2019), the fixation of a deleterious mutation by genetic drift can generate a similar selective sweep effect to that caused by the fixation of a beneficial mutation, for mutations with the same magnitude of selection coefficient. Moreover, Tajima (1990) demonstrated that, on average, there is a ~42% mean reduction in diversity at a site where a neutral mutation has recently become fixed by genetic drift. While the mean time to fixation for this class of mutation in diploids is well known to be 4*N_e_* generations (Kimura & Ohta 1969), this is associated with a wide variance of approximately 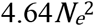 (Kimura 1970), such that neutral mutations may also fix relatively rapidly (in less than *N_e_* generations) and generate an appreciable but highly localized sweep effect (see Tables 1 and 2 of Tajima 1990).

**Table 1:**
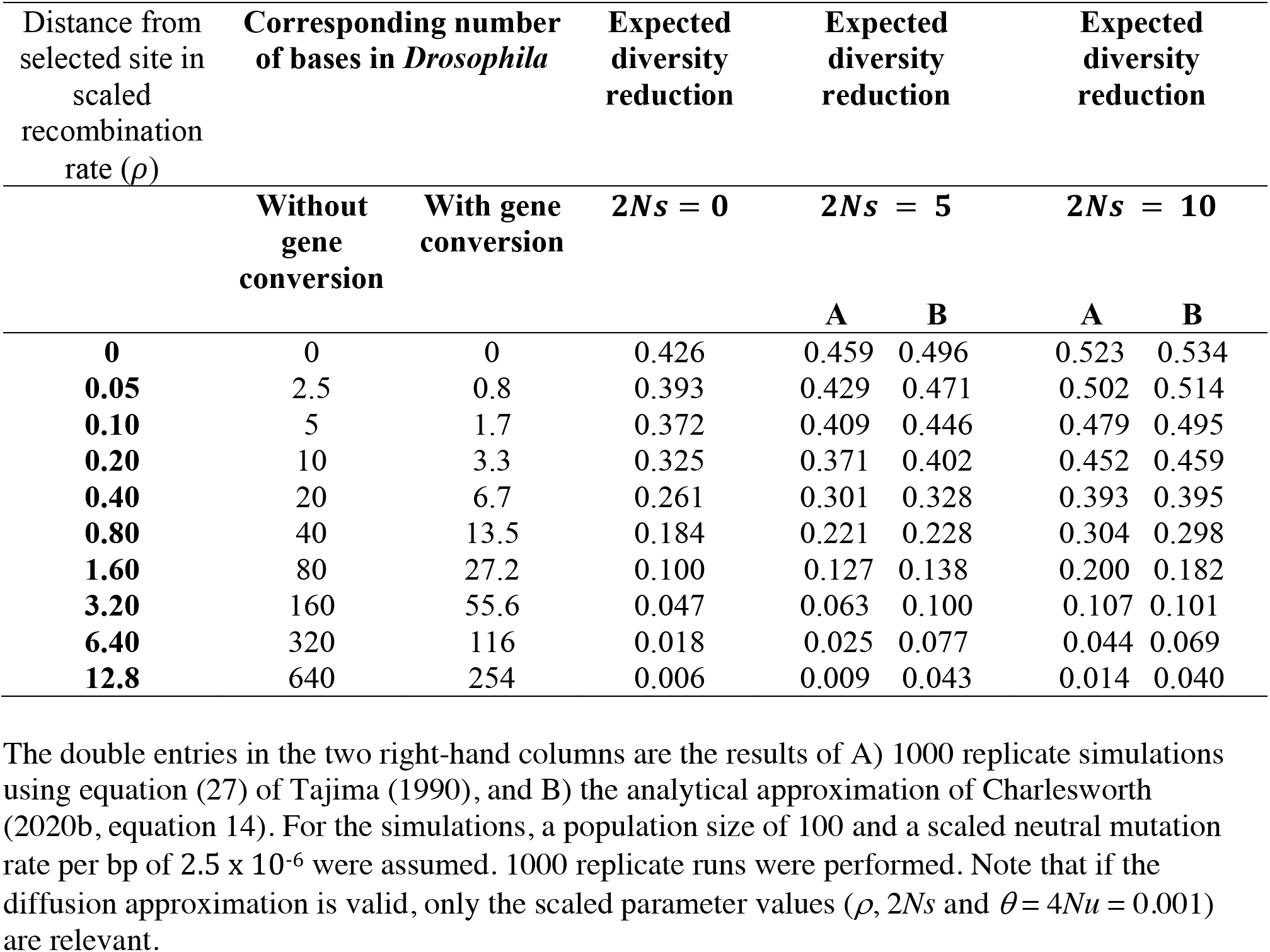
Reduction in nucleotide diversity at linked neutral sites relative to the neutral value after the fixation of a weakly deleterious or advantageous semi-dominant mutation. The distance between the selected and neutral site is presented in units of the scaled recombination rate (*ρ*), as well as the number of bases corresponding to *Drosophila*-like populations.

**Table 2:** Mean genome-wide and outlier *F_ST_* values (calculated in windows of 500 bp (labeled ‘Constant total sites’), or 10 SNPs (labeled ‘Constant SNPs’)).

| Evolutionary model | Mean $F_{ST}$ | | Outlier $F_{ST}(1\%)$ | | Outlier $F_{ST}(1\%)/$<br>Mean $F_{ST}$ | |
| --- | --- | --- | --- | --- | --- | --- |
|  | Constant total sites | Constant SNPs | Constant total sites | Constant SNPs | Constant total sites | Constant SNPs |
| Neutrality | 0.0374 | 0.0363 | 0.0874 | 0.1628 | 2.3369 | 4.4838 |
| Purifying selection | 0.0365 | 0.0352 | 0.0880 | 0.1454 | 2.4110 | 4.1296 |
| Purifying selection (rescaled DFE) | 0.0666 | 0.0613 | 0.1585 | 0.2637 | 2.3799 | 4.2999 |
| With 1% weak positive selection | 0.0378 | 0.0362 | 0.1443 | 0.1631 | 3.8143 | 4.5057 |
| With 1% strong positive selection | 0.2353 | 0.2347 | 0.9108 | 0.6689 | 3.8705 | 2.8507 |
| Neutral values, with the Li & Stephan (2006) bottleneck | 0.2828 | 0.2557 | 0.4154 | 0.6956 | 1.4689 | 2.7204 |

Of course, the probabilities of fixation of deleterious and beneficial fixations differ greatly. However, given that the input of deleterious mutations is much higher than the input of beneficial mutations each generation (see reviews by Eyre-Walker & Keightley 2007; Bank *et al.* 2014a), the potential contribution of such deleterious sweeps to levels and patterns of nucleotide variation, as well as divergence between populations and species, remains an important open question (Charlesworth 2020a). An alternative way of viewing this issue, as discussed by Gillespie (1994) and Charlesworth & Eyre-Walker (2007), is that under a model of constant selection and reversible mutation between two alternative nucleotide variants, statistical equilibrium with respect to the frequencies of sites fixed for the alternatives implies equal rates of beneficial and deleterious substitutions per unit time. It is important to note, however, that only deleterious mutations with selection coefficients on the order of the reciprocal of the population size have significant probabilities of fixation (Fisher 1930; Kimura 1964), implying that the substitutions concerned involve only very weakly selected mutations. The effects on diversity statistics of sweeps of very weakly selected mutations, including those of deleterious mutations, appears to have been investigated previously only by Mafessoni & Lachmann (2015).

A starting point for investigating this problem is the distribution of fitness effects of new mutations (the DFE). There is substantial evidence from both empirical and experimental studies that the DFE of new mutations is bimodal - consisting of a strongly deleterious mode, and a weakly deleterious / neutral mode that may contain a beneficial tail under certain conditions (*e.g.*, Crow 1993; Lynch *et al.* 1999; Sanjuán 2010; Jacquier *et al.* 2013; Bank *et al.* 2014b). While the calculations and simulations presented below represent a general approach to addressing this topic, we have necessarily chosen a specific DFE realization and species for illustration. Specifically, Johri *et al.* (2020) recently presented an approximate Bayesian (ABC) approach, which represents the first joint estimator of the DFE shape together with population history, and which corrects for the effects of background selection (BGS; Charlesworth *et al.* 1993). They estimated that a substantial proportion of new mutations in coding regions have mildly deleterious effects on fitness, emphasizing the importance of further understanding the consequences of such mutations in dictating observed polymorphism and divergence. Furthermore, they found it unnecessary to invoke a beneficial mutational class in order to fit the data from the African population of *Drosophila melanogaster* that they considered.

This analysis provides a basis for exploring the possible implications of sweeps of deleterious mutations, a topic that has been neglected. Here, we re-examine this question, considering both single and recurrent substitution models, which we use to examine the possibility that genomic scans for positive selection may in fact also be identifying deleterious fixations, when based on: a) levels and patterns of variation; b) population differentiation; and c) species-level divergence. Our results suggest that, while this phenomenon is unlikely to be a major factor in polymorphism-based scans within populations, it may be a serious confounder in among-population-based analyses.

## METHODS

### Analytical calculations

For convenience, we assume a Wright-Fisher population of *N* randomly mating diploid individuals throughout the analytical section. This assumption of an absence of population structure is common and reasonably well-justified in organisms with low *F_ST_* like *D. melanogaster*, although the investigation of these dynamics in the presence of structure would be of great value in the future. Further, the selection coefficient for homozygotes is constant over time and is denoted by *s*, with *s_a_* > 0 and *s_d_* > 0 representing selection for and against homozygotes, respectively. In addition, the DFE for semi-dominant deleterious mutations is assumed to be discrete, with four fixed classes of mutations given by 0 <2*Ns_d_* ≤ 1, 1 <*Ns*_*d*_ ≤ 10, 10 <2*Ns_d_*≤ 100 and 100< 2*Ns_d_* ≤ 2*N*, where mutations are assumed to follow a uniform distribution within each class. These assumptions concerning the distribution of *s_d_* were made in order to simplify integration over the DFE (Johri *et al*. 2020; and see Johri *et al.* 2021).

#### Probability of fixation

The fixation probability (*P_fix_*) of a new semi-dominant mutation with an initial frequency of 1/(2*N*) in a Wright-Fisher population of size *N* was calculated using Equation 10 of Kimura (1962):

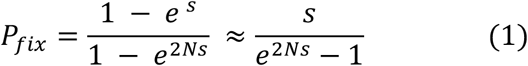

Note that this equation assumes demographic equilibrium and independence between the selected sites (violations of this assumption are investigated below), as well as |*s*| << 1.

#### Contributions to divergence

Under the DFE model described above, the number of fixations *N_fix_* (*i*) expected per generation per site for a given DFE class *i* of deleterious mutations is given by the following expression:

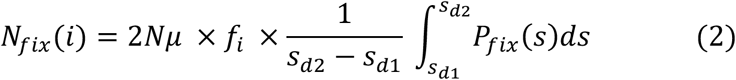

where *μ* is the total mutation rate per site per generation, *f*_i_ represents the proportion of new mutations belonging to the *i*^th^ DFE class and *s*_*d*2_ and *s*_*d*1_ represent the upper and lower bounds to the DFE class, respectively. The integral in equation (2) was evaluated analytically by means of an infinite series representation (derivation provided in Appendix 1), which was validated using the “integrate” function in R (R Core Team 2018).

#### Probability of fixation when correcting for the effects of background selection (BGS)

The fixation probability (*P_fixB_*) of a new semi-dominant mutation with the homozygous selection coefficient *s* was calculated as:

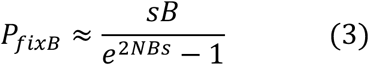

where *B* = *N_e_*/*N* represents the effective population size in the presence of BGS, calculated from the corresponding ratio of neutral diversity with and without BGS (see, for example, Campos & Charlesworth 2019). The mean probability of fixation over an interval of selection coefficients, assuming that mutations are uniformly distributed in this interval (*P_fixB_*(*γ*_1_ − *γ*_2_)), was calculated as:

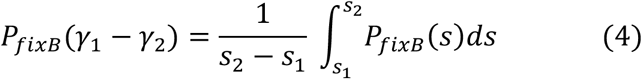

where *γ*_1_ = 2*NBs*_1_, *γ*_2_ = 2*NBs*_2_ and *γ*_2_ and *γ*_1_ represent the upper and lower bound to the interval, respectively.

#### Waiting and fixation times

The waiting time (*t_w_*) between fixations under a Poisson process was calculated as follows:

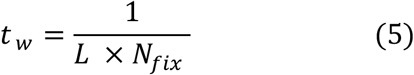

where *L* represents the number of functional sites under consideration.

In order to compare the results of simulations to theoretical expectations, the expected time to fixation of a semi-dominant mutation was also calculated by numerically integrating equation (17) of Kimura & Ohta (1969), using Simpson’s rule (Atkinson 1989).

#### Reduction in diversity due to a single sweep

The expected reduction in pairwise nucleotide site diversity at the end of a sweep (–Δ*π*), relative to the diversity in the absence of selection, was calculated using equation (14a) of Charlesworth (2020b) for a non-zero rate of recombination (omitting a factor that describes the effect of background selection):

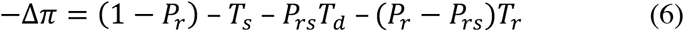

where *T_s_* is the mean coalescent time (in units of *N*) for a pair of alleles sampled at the end of a sweep, *T_d_* is the duration of the deterministic phase of the sweep (*i.e*., excluding the initial and final stochastic phases) in units of *N*, *T_r_* is the mean time to a recombination event during the sweep, conditioned on the occurrence of a recombination event; *P_r_* is the probability of at least one recombination event transferring a sampled allele onto the wild-type background during the sweep, and *P_rs_* is the probability that there is only a single such recombination event. For large values of the ratio of the rate of recombination between the neutral and selected site (*r*) to the magnitude of the selection coefficient, this expression can become negative, in which case it is reset to zero.

We also used simulations based on equations (27) of Tajima (1990), which provide recursion equations for the expectation of the pairwise diversity at a neutral locus linked to a selected locus, conditional on a given trajectory of allele frequency change at the selected locus, as described by Charlesworth (2020b). These equations require only the validity of the diffusion equation approximation. Binomial sampling of allele frequencies post-selection in each generation was used to generate the trajectories of change at the selected locus, using the standard selection equation for a single locus to calculate the deterministic change in allele frequency each generation. Application of the recursion equations to a trajectory of allele-frequency change simulated in this way gives one realization of Δ*π*, and the overall expected value of Δ*π* can be obtained from the mean of the simulated values over a large number of replicates. It was found that 1000 replicates gave very accurate estimates of Δ*π*, with ratios of the standard errors to the means of < 5% for the parameter sets used here. Due to the symmetry in the sojourn time (conditional on fixation) in a given interval of allele frequency for beneficial and deleterious mutations when semi-dominance is assumed (Maruyama and Kimura 1974), trajectories for weakly deleterious mutations of selection coefficient *s_d_* were simulated assuming a beneficial selection coefficient *s_a_*. Note that this symmetry does not apply to partially dominant or partially recessive mutations – see Ewens (2004, pp. 170-171) for a full discussion.

#### Population genetic parameters used for analytical calculations

The parameters for the calculations were chosen to match those estimated from *D. melanogaster* populations, estimated from exonic sites. Mutations occurred at a rate *μ* per basepair (bp) per generation, and were assumed to be a mixture of neutral, nearly neutral and weakly deleterious mutations. For this analysis, *μ* = 3 × 10^−9^ (Keightley *et al.* 2009, 2014). The sex-averaged rate of crossing over per bp (*r_c_*) was assumed to be equal to 10^−8^ per generation, the mean value for *D. melanogaster* autosomes (Fiston-Lavier *et al.* 2010), and the effective population size was 10^6^ (Arguello *et al.* 2019; Johri *et al.* 2020), and 10 generations per year were assumed. Given estimates of neutral divergence between *D. melanogaster* and *D. simulans*, this means that *t* ~21.3 × 10^6^ generations elapsed since their common ancestor (Li *et al.* 1999; Halligan & Keightley 2006), corresponding to 2.13 million years.

Because non-crossover associated gene conversion is an important source of recombination between closely linked sites in *Drosophila*, it was assumed to occur uniformly across the genome, independently of local differences in the rate of crossing over, as indicated by the data on *D, melanogaster* (Comeron *et al.* 2012; Miller *et al.* 2016). The sex-averaged rate of initiation per bp of conversion events was *r_g_* = 10^−8^ per bp per generation, and there was an exponential distribution of tract length with a mean of *d_g_* = 440 bp (Comeron *et al.* 2012; Miller *et al.* 2016). The net rate of recombination between sites separated by *z* bp, *r*(*z*), is the sum of the contributions from crossing over and non-crossover gene conversion, given by the formula of Frisse *et al.* (2001):

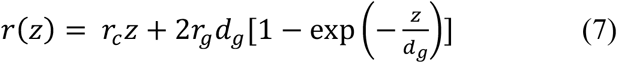

For some of the results presented below, only the net rate of recombination *r*, or its value scaled by the effective population size, *ρ* = 2*Nr*, was used. In the presence of gene conversion with the parameters described above, the corresponding value of *z* can then be obtained by Newton-Raphson iteration of the equation *r*(*z*) − *r* = 0, assuming that the rate of gene conversion does not vary with the rate of crossing over,

### Simulations

#### Simulating individual fixations in a single population

SLiM 3.3.1 (Haller & Messer 2019) was used to simulate a genomic element of length 10 kb. For the single sweep model, in order to quantify the hitchhiking effect of a single fixation, selection only acted on a single site in the middle of the region, with all other sites evolving neutrally. Simulations were performed for five different values of the scaled selection coefficient: 2*Ns* = 0, −1, +1, −5, and +5. Population genetic parameters resembling those of *D. melanogaster* were utilized (as defined above) for illustrative purposes. In order to perform simulations efficiently, population size was scaled down by a factor of 100 while the mutation and recombination rates were correspondingly scaled up by the same factor. Simulations were run for a burn-in period of 10^5^ generations (10*N*_sim_) after which a mutation with a scaled selection coefficient of 2*Ns* was introduced at the selected site. Simulations in which the introduced mutation reached fixation were retained for analysis. Fifty diploid individuals were sampled at the completion of the simulations to mimic generally-available population-genomic data, and population genetic summary statistics were calculated using *Pylibseq* (Thornton 2003).

#### Simulating fixations in a single population in the presence of other deleterious mutations

SLiM 3.1 (Haller & Messer 2019) was used to perform simulations resembling both *Drosophila*-like and human-like populations in order to assess the effects of Hill-Robertson interference (Hill & Robertson 1966) amongst deleterious mutations (reviewed by Charlesworth 2012). For parameters resembling the *D. melanogaster* populations, a ~15kb region which was composed of 2 genes, each with 5 exons (of length 300 bp) and 4 introns (of length 100 bp), and with intergenic lengths of 4kb was simulated. Intergenic and intronic regions experienced only effectively neutral mutations, while the exons experienced mutations with the DFE inferred by Johri *et al.* (2020). The population parameters were chosen to mimic the Zimbabwe population of *D. melanogaster*, with *N_e_* = 1.95 × 10^6^, mutation rate = 3.0 × 10^−9^ per site/generation and recombination rate = 1 × 10^−8^ per site/generation (Arguello *et al.* 2019).

For parameters resembling human populations, a ~60kb region composed of 2 genes, each with 5 exons (of length 300 bp) and 4 introns (of length 2 kb), and with intergenic lengths of 15kb was simulated. Note that these values are nearer to the median than the mean values of the distribution of lengths of genomic regions in humans, and provide a more conservative assessment of potential interference effects. Intergenic and intronic regions experienced only neutral mutations, while the exons experienced mutations with the DFE inferred by Huber *et al.* (2017), such that *f*_0_ = 0.51, *f*_1_ = 0.14, *f*_2_ = 0.14 and *f*_3_ = 0.21. The population parameters used were *N_e_* = 10^4^, mutation rate = 1.2 × 10^−8^ per site/generation and recombination rate = 1 × 10^−8^ per site/generation.

Simulations were conducted with *N* = 10,000 diploid individuals for both species, and mutation and recombination rates were scaled appropriately (by the factor 195 for *D. melanogaster* and 1 for humans). In order to estimate the rates of fixation of deleterious mutations and their effects on linked sites, the simulations were run for 100*N* generations (of which 10*N* generations represent the burn-in period) and were replicated 100 times for *D. melanogaster* and 600 times for humans. Our simulations resembling *D. melanogaster* populations resulted in a mean *K_A_/K_S_* value of 0.08, which is consistent with an observed mean autosomal value of 0.145 for a comparison with a closely related species (Campos *et al.* 2014), where 50% of substitutions are expected to be due to beneficial mutations.

The total number of observed substitutions (presented in Supp Table 1) were used to obtain the frequencies of fixation and times to fixation of deleterious mutations, and nucleotide site diversities at linked neutral sites immediately post-fixation, in order to quantify the potential effects of interference among selected mutations. The frequency of fixations of weakly deleterious mutations with scaled selection coefficients distributed between *γ*_1_ and *γ*_2_, *P_fixsim_*(*γ*_1_ − *γ*_2_),)) where 1 ≤ *γ*_1_ < *γ*_2_ ≤ 10 (*i.e*., for a set of mutations that are a subset of the weakly deleterious class of mutations) was calculated as:

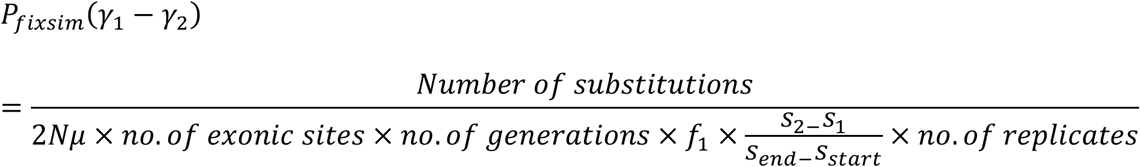

where 2*Ns*_*start*_ = 1, 2*Ns*_*end*_ = 10; and *γ*_1_ = 2*Ns*_1_, *γ*_2_ = 2*Ns*_2_ when not correcting for BGS, and *γ*_1_ = 2*NBs*_1_, *γ*_2_ = 2*NBs*_2_ when correcting for BGS. Again, *B* is the ratio of the expected nucleotide diversity at a neutral site in the presence of BGS to its value in the absence of BGS.

#### Simulating multiple populations and FST-based analyses

Based on a recently inferred demographic history of African and European populations of *D. melanogaster* using populations sampled from Beijing, the Netherlands, and Zimbabwe (Arguello *et al.* 2019), an ancestral population of size of 1.95 × 10^6^ was simulated, which split into two populations (6.62 × 10^4^ generations ago) of constant size: 3.91 × 10^6^ and 4.73 × 10^5^, representing African and European populations, respectively. Note that, although Arguello *et al.* inferred recent growth in both populations, we have assumed constant sizes in order to avoid confounding effects of such growth on the fixation probabilities of mutations, and we also assumed no migration between the two populations. For scenarios following the demographic model presented in Arguello *et al.* (2019), simulations were also performed for the extreme values of the 95% CIs for the current size of the Zimbabwe population (3.02 × 10^5^, 4.69×10^6^), the current size of the European population (2.03 × 10^5^, 3.89×10^6^), and the time of split (1.17×10^4^, 1.03×10^5^). Ten replicates of each combination of the above-mentioned values (*i.e*., 8 different combinations) were simulated in order to account for uncertainties in parameter estimates. For the purpose of *F_ST_* analyses, a functional region of size 10 kb was simulated, in which mutations had selection coefficients given by the DFE inferred in Johri *et al.* (2020). Specifically, the DFE was given by a discrete distribution of four fixed bins with 24.7% of mutations belonging to the effectively neutral class (*f*_0_: 0 ≤ 2*Ns_d_* <1), 49.4% to the weakly deleterious class (*f*_1_: 1 ≤ 2*Ns_d_*< 10), 3.9% to the moderately deleterious class (*f*_2_: 10 ≤ 2*Ns_d_* < 100), and 21.9% to the strongly deleterious class of mutations (*f*_3_: 100 < 2*Ns_d_* ≤ 2*N*).

In this two-population framework, four separate scenarios were tested: 1) the DFE remained scaled to the ancestral population size (*i.e*., the distribution of selection coefficients remained constant throughout, making selection effectively weaker in the smaller derived (European) population); 2) the DFE was rescaled after the population split with respect to subpopulation-specific sizes (*i.e*., both populations experienced equal proportions of mutations belonging to each DFE class as defined in terms of 2*Ns_d_*), such that selection was equally strong in both populations - this is an arbitrary biological model that is simply chosen for comparison, as one would naturally expect selection to be effectively weaker in the derived population as in scenario 1; 3) in addition to this neutral and deleterious DFE, 1% of all mutations were mildly beneficial with selective effects drawn uniformly from the interval 1 ≤ 2*Ns_a_* ≤ 10, where *s*_*a*_ is the increase in fitness of the mutant homozygote; and 4) in addition to this neutral and deleterious DFE, 1% of all mutations were strongly beneficial with 2*Ns*_*a*_ = 1000. In scenarios 3 and 4, the deleterious and beneficial DFEs were scaled to the respective sizes of each population post-split, so that the proportions of beneficial mutations were the same in the two populations. Note that we refer to selection as being weak when 2*Ns* is less than 10. In order to assess the role of population bottlenecks in generating neutral outliers, a fifth scenario was simulated in which there was no selection and the demographic history was that inferred by Li & Stephan (2006) - a model that involves a much larger size reduction than inferred in the Arguello *et al.* (2019) model utilized above. The parameters of both demographic models are provided in Supp Table 2.

Fifty diploid individuals were sampled from both populations in order to calculate *F_ST_*. All sites that would be considered polymorphic in the metapopulation were used to calculate *F_ST_* (*i.e*., sites fixed either in one population or both populations (for different alleles) were also included in *F_ST_* calculations). *F_ST_* was calculated in sliding windows across the genomic region for: a) windows containing a constant number of SNPs (10 SNPs) using the package PopGenome (Pfeifer *et al.* 2014) in R, and b) for windows representing the same total number of bases (500 bp) using *Pylibseq* 0.2.3 (Thornton 2003). *F_ST_* was calculated for both cases by the method of Hudson *et al.* (1992). *F_ST_* was also calculated individually for different mutation types (*i.e*., for neutral, weakly deleterious and beneficial mutations, by simply restricting the calculations to segregating sites of the specific mutation type). Although there will be an upper bound to the *F_ST_* values obtained in this way, which is determined by the frequency of the most frequent allele in the metapopulation (Jakobsson *et al.* 2013), the detection of outliers should not be affected by this procedure and should mimic the empirical practice. In order to evaluate the performance of haplotype-based population differentiation statistics, *φ_ST_* was calculated following Excoffier *et al.* (1992), by performing the Analysis of Molecular Variance (AMOVA) approach implemented in the *R* package *Pegas* (http://ape-package.ird.fr/pegas.htlm) on SNPs present in 500 bp, 1 kb and 2 kb sliding windows.

#### Simulating multiple species, and divergence-based analyses

McDonald-Kreitman (MK) tests (McDonald & Kreitman 1991) were performed to investigate the degree to which substitutions of mildly deleterious mutations might affect the inference of divergence due to positive selection. A population resembling a *D. melanogaster* African population was simulated under demographic equilibrium. Ten independent replicates of a 10 kb protein-coding region were considered such that every third position was neutral (representing synonymous sites) while all other sites represented nonsynonymous positions. Nonsynonymous sites experienced purifying selection given by a DFE that comprised only the non-neutral bins (*i.e., f*_1_, *f*_2_, and *f*_3_, in the same proportions described above), while the neutral sites experienced mutations that belonged to class *f*_0_. For comparison, simulations were performed in which 10% or 20% of nonsynonymous mutations were also neutral. In all cases, the simulation was run for 20*N* generations (where *N* = 10^4^). The number of segregating nonsynonymous (*P_N_*) and synonymous (*P_S_*) sites were estimated by sampling 50 diploid individuals from the population from the functional and neutral regions respectively. The number of fixed substitutions occurring at the functional sites (*D_N_*) and neutral sites (*D_S_*) were calculated post burn-in (*i.e.*, after 10*N* generations), and then rescaled to the number of generations since the *D. melanogaster* ancestor.

In order to correct for mildly deleterious mutations segregating in populations, the proportion of adaptive substitutions (*α*) was also inferred by implementing a variant of the test referred to as the asymptotic MK test. Messer & Petrov (2013) suggested plotting the derived allele frequency (*x*) of variants against *α* inferred using the number of segregating sites (*P(x)*) at that derived allele frequency (*i.e*., 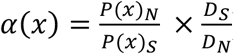), and showed that the asymptote of this curve would tend towards the true value of *α*. The asymptotic MK test was performed using a web-based tool available at: http://benhaller.com/messerlab/asymptoticMK.html (Haller & Messer 2017). For the purpose of the asymptotic MK test, values of *P_N_*/*P_S_* were binned with a bin size of 0.05, and the curve-fitting (of *α*(*x*) with respect to *x*) was restricted to derived allele frequencies between 0.1-1.0

##### Data Availability Statement

All scripts used in this study are provided in the Github repository: https://github.com/paruljohri/Deleterious_Sweeps.

## RESULTS & DISCUSSION

### Theoretical expectations

As has been long appreciated (Fisher 1930; Wright 1931), equation (1) implies that the probability of fixation of a mutation in the strongly deleterious DFE classes is vanishingly small. However, the weakly deleterious class may contribute substantially to divergence (Figure 1a). We derived an analytical approximation for the probability of fixation for mutations with a DFE represented as a combination of four non-overlapping uniform distributions (see Methods and Appendix 1). In a Wright-Fisher population, the ratio of the mean probability of fixation of mutations with fitness effects between 1 < 2*Ns_d_* ≤ 5, relative to the probability of fixation of effectively neutral mutations (0 < 2*Ns_d_* ≤ 1), is 0.27 (note that the value of *N* is irrelevant if the diffusion approximation holds). This ratio rapidly declines to 0.01 for mutations with fitness effects 5 < 2*Ns_d_* ≤ 10 (Figure 1a).

**Figure 1:**
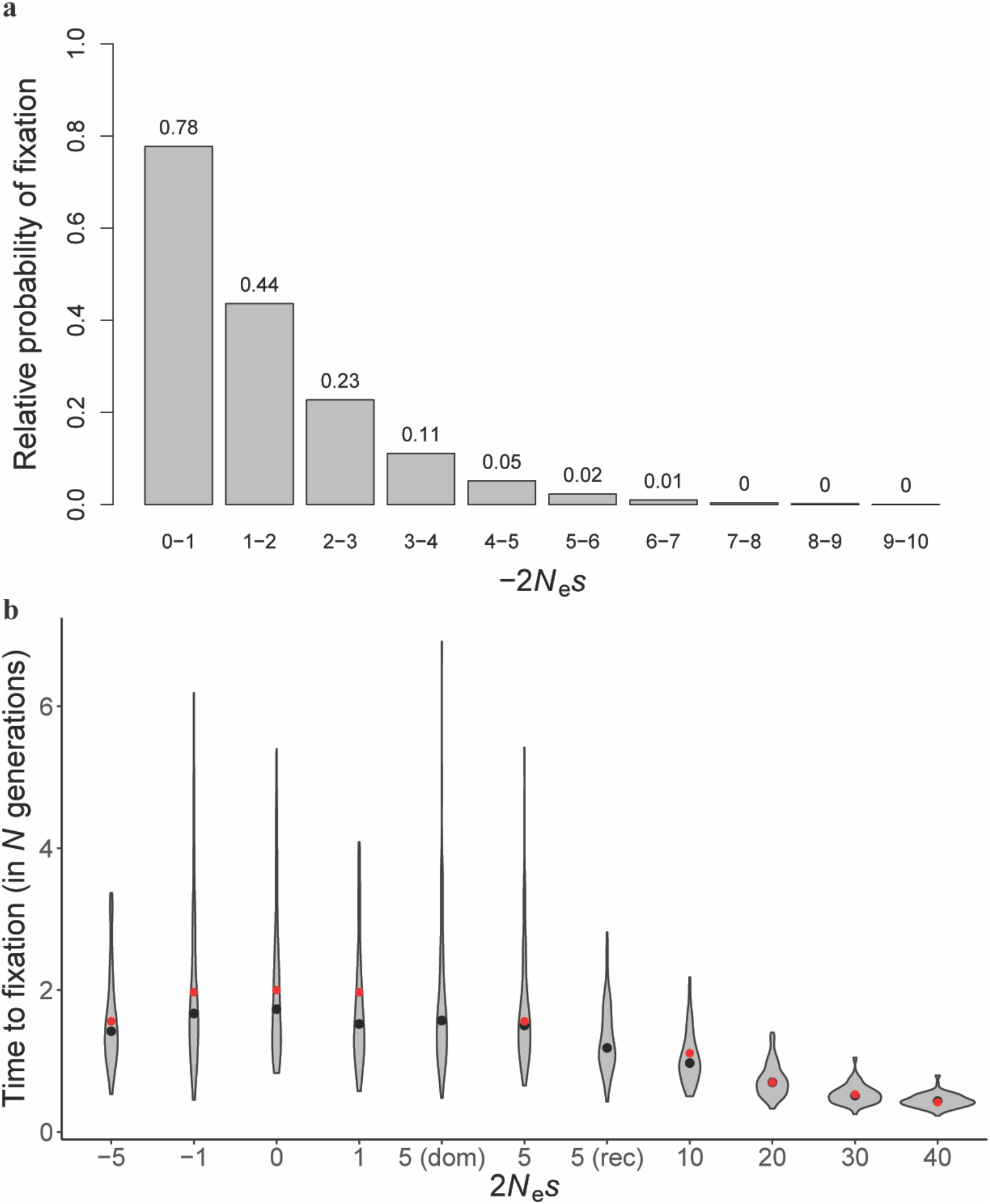
(a) Frequencies of fixations of weakly deleterious mutations relative to those of neutral mutations. (b) The distribution of fixation times (conditional on fixation) of mutations with varying selective effects, obtained from 100 simulated replicates. Fixation times are measured as the time taken for the mutant allele to spread from frequency 1/(2*N*) to frequency 1. Black solid circles are the means of the distributions obtained from simulations, and red solid circles are the mean expectations obtained by numerically integrating the expression of Kimura & Ohta (1969). The dominance coefficient is 0.5 for all mutations except in the cases “dom” and “rec” where *h* = 1 and 0, respectively.

Taking the recently estimated DFE for the Zambian *D. melanogaster* population (Johri *et al.*, 2020) as an example, a question of interest concerns the probabilities of fixation of different classes of mutations and their contributions to population- and species-level divergence (Supp Figure 1). In the absence of positive selection, the weakly deleterious class of mutations would be expected to contribute 7.2% of the total divergence in exonic regions, while effectively neutral mutations would be expected to contribute 92.8%. If we were to assume that approximately 50% of all substitutions in *Drosophila* have been fixed by positive selection (Eyre-Walker and Keightley 2009; Campos *et al.* 2017), weakly deleterious mutations are still likely to have contributed 3.5% of the total divergence in functional regions and possibly much more in regions experiencing reduced selective constraints.

### Hitchhiking effects of deleterious sweeps: levels and patterns of variation

We next considered the fixation times of these mutations contributing to divergence, as well as the expected waiting time between fixations. Using population parameters relevant for *D. melanogaster* (see Methods), the neutral and deleterious DFE of Johri *et al.* (2020), and assuming that 60% of the *D. melanogaster* genome (of size 140 Mb) is functional, equation (5) shows that the genome-wide waiting time between successive weakly deleterious fixations is ~ 83 generations. Hence, the ~2.13 million years estimated for the time since the *D. melanogaster* and *D. simulans* split is expected to equate to many such fixations.

As expected, there is no significant difference (*p* = 0.16; Student’s *t*-test) in fixation times between the mildly beneficial (3.2*N* generations; SD: 1.5*N*) *vs* mildly deleterious mutations (2.9*N* generations; SD: 1.1*N*) with the same fitness effects (2*Ns*_*a*_=2*Ns_d_*=5) (Supp Table 3). Importantly however, the variance in fixation times of weakly selected mutations is extremely large, such that the faster tails of the fixation time distributions for both 2*Ns_d_* = 5 and 2*Ns* = 0 occupy ~*N* generations, which corresponds to a sweep effect of the same size as the mean for 2*Ns*_*a*_ = 30 (Figure 1b). In other words, weakly deleterious and neutral fixations can match the sweep effects of comparatively strongly selected beneficial fixations.

We evaluated the impact of these fixations on observed genomic variation in two ways. The first corresponds to a model of a single sweep event, which is relevant to the literature on detecting signatures of individual fixations, as done in genomic scans for positively selected loci (*e.g*., Harr *et al.* 2002; Glinka *et al.* 2003; Haddrill *et al.* 2005; Nielsen *et al.* 2005; see also Jensen 2009; Stephan 2019). Such scans operate under the assumption that the selective sweeps in question have achieved fixation immediately prior to sampling, due to the rapid loss of signal as the time since fixation increases (Kim & Stephan 2002; Przeworski 2002). The second corresponds to a recurrent substitution model, which is relevant to the recurrent sweep literature that attempts to quantify the effect of selective sweeps on genome-wide patterns of variability and their relation to rates of recombination (*e.g*., Wiehe & Stephan 1993; Kim 2006; Andolfatto 2007; Macpherson *et al.* 2007; Jensen *et al.* 2008; Campos & Charlesworth 2019; Charlesworth 2020b; see review by Sella *et al.* 2009).

The reduction of diversity resulting from the fixation of a mildly deleterious or beneficial semi-dominant mutation with a very small selective effect is not expected to be substantially different from that caused by the fixation of a selectively neutral mutation (Tajima 1990). The widely-used approximation of Barton (2000) for the reduction in diversity relative to neutrality caused by the sweep of a semi-dominant mutation, Δ*π*= (2*Ns_a_*)^(−4*r*/*s*)^, suggests that, when 2*Ns*_*a*_ = 5, diversity will only be reduced by more than 20% in a region for which *r*/*s* ≤ 0.2, corresponding to approximately 20 bp for a typical *D. melanogaster* recombination rate of 3 × 10^−8^ per bp, including the contribution from gene conversion as given by equation (7), and assuming an effective population size of ≥ 10^6^.

However, this formula assumes that fixations are so fast that swept alleles that have failed to recombine onto a wild-type background experience no coalescent events during the duration of the sweep (Hartfield & Bataillon 2020; Charlesworth 2020b), which is unlikely to be true for weakly selected mutations, such that NIsI is close to 1. We therefore used equation (6) for analytical predictions (Figure 2, Table 1, Supp Table 4), which is based on the results of Charlesworth (2020b). But this equation is also likely to be inaccurate with weak selection, because it assumes that the trajectory of allele frequency change is close to that for the deterministic case, except for the initial and final stochastic phases at the two extremes of allele frequencies. We therefore also used simulations based on equations (27) of Tajima (1990) to predict sweep effects, as described in the Methods.

**Figure 2:**
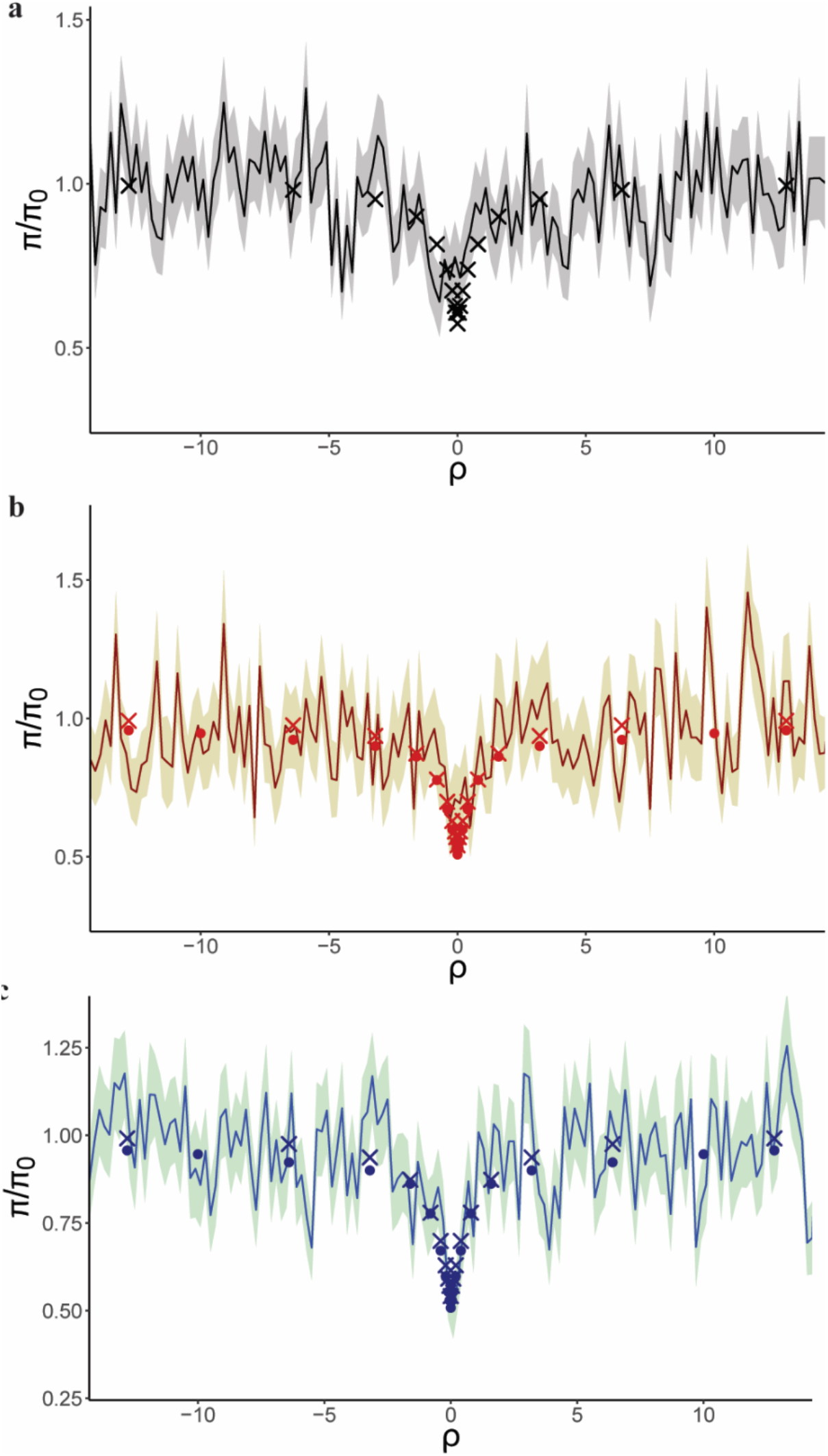
Recovery of nucleotide diversity per site (*π*) relative to the mean value in the absence of selection neutrality (*π*_0_), around a recent fixation (shown at position 0 on the x-axis). The target site has experienced (a) a neutral fixation (2*Ns* = 0; black lines), (b) a weakly deleterious fixation (2*Ns_d_*= 5; red line), and (c) a weakly beneficial fixation (2*Ns*_*a*_ = 5; blue line). Solid lines represent mean values of 100 replicates, shaded regions correspond to 1 SE above and 1 SE below the mean. Solid circles show the theoretical predictions using equation (14) of Charlesworth 2020b; and crosses correspond to simulations based on equation (27) of Tajima (1990).

With the full simulations of a 10 kb region, in which we assumed a population size of 10^6^ and mutation rate of 3 × 10^−9^ /site/generation, with 2*Ns*_*a*_= 5 the nucleotide diversity 10 bp (~*ρ* = 0.2) around the selected site (*i.e*., 5 bp in both directions) was 0.0058 (SE: 0.0012), corresponding to a reduction of 52% below the neutral value. For 2*Ns* = 10, the 10-bp nucleotide diversity was 0.0083 (SE: 0.0017), corresponding to a reduction of 31%. The observed reduction in both cases almost fully recovers to the expected level under neutrality within 500 bp (*ρ* ≈ 10; Figure 2). A similar pattern is seen in Table 1 and Supp Table 4, which shows both the analytical predictions and those based on Tajima’s equations, which agree surprisingly well except at the two highest rates of recombination displayed. For comparison, the results of simulating neutral fixations are also shown in Table 1 and Figure 2. The case with *Ns* = 10 is at the higher limit of what is likely to be produced by sweeps of deleterious mutations, given that the ratio of the fixation probability given by equation (1) to the neutral value of 1/(2*N*) is then approximately 0.0045, compared to 0.034 with *Ns* = 5.

Mafessoni & Lachmann (2015) showed that that fixations of weakly selected, highly dominant favorable mutations - or highly recessive deleterious mutations - could reduce diversity at linked sites by a *smaller* amount than fixations of neutral mutations (with no recombination, these are associated with a diversity reduction of 42% below the mean neutral value), with a maximum effect when 2*Ns* is approximately 2 (and *h* = 1 for favorable mutations, and 0 for deleterious ones). We have confirmed this unexpected observation using Tajima algorithm simulations (Supp Table 5), finding that it exists even for 2*Ns* = 5 (Supp Figure 2; Supp Table 5). However, our use of *h* = 0.5 and 2*Ns* ≥ 2.5 means that this phenomenon is absent from the results presented above.

Because gene conversion is an important contributor to recombination between closely linked sites (Miller *et al.* 2016), the effects of fixations in the presence of gene conversion are restricted to a region that is about one-third of the distance in the absence of gene conversion (Table 1). Thus, a greater than 20% reduction in nucleotide diversity for 2*Ns* = 5 [2*Ns* = 10] was observed up to *ρ* = 0.8 [*ρ* = 1.6], corresponding to 14 bp [27 bp] with gene conversion, and 40 bp [80 bp] without gene conversion. This quite localized effect is similar for weakly beneficial, weakly deleterious, and neutral mutations.

Given the relatively faster mean speed (3.1*N* generations) of fixation of weakly selected semi-dominant mutations compared to the neutral expectation (4*N* generations), they should also result in small distortions of the SFS at closely linked neutral sites. We observed a slight skew towards rare variants (as measured by Tajima’s *D*, Supp Table 6) restricted to ~50 base pairs from the selected site immediately after fixation (for 2*Ns_d_* = 2*Ns*_*a*_ = 5). This highly localized distortion of the SFS is probably too weak to play any important role in generating false positives in genomic scans. Indeed, owing to the inherent stochasticity involved in the underlying processes, such scans generally only have power to detect very strongly selected fixations (often requiring values of 2*Ns*_*a*_ > 1000 in order to observe appreciable true positive rates: Crisci *et al.* 2013). Viewed in another way, given that the false-positive rates associated with genomic scans may often be inflated well above true-positive rates owing to the underlying demographic history of the population (*e.g.*, Teshima *et al.* 2006; Thornton & Jensen 2007; Crisci *et al.* 2013; Harris *et al.* 2018), demography is probably a much stronger confounder than deleterious sweeps in polymorphism-based scans.

Finally, using a recurrent fixation model, we have examined the steady-state impact of weakly deleterious sweeps (Figure 3). Here we used a model of a gene with five exons of 100 codons each, with 70% of exonic mutations subject to selection, which were separated from each other by 100 bp introns, as described by Campos & Charlesworth (2019) and Charlesworth (2020b). Five equally large classes of sites subject to deleterious mutations were modeled, with the lowest scaled selection coefficient being 2*Ns_d_* = 2.5 and the largest 2*Ns_d_* = 10; as described above, a uniform distribution of 2*Ns_d_* values was assumed within each class. The theoretical predictions used the method for predicting the effects of recurrent sweeps of Charlesworth (2020b), based on Equation 6 above. For consistency with the simulations of population divergence described below, the population size was assumed to be 1.95 × 10^6^, giving a neutral diversity of 0.0234 with a mutation rate of 3 × 10^−9^. Substitution rates of deleterious nonsynonymous mutations were calculated from equations (1) and (2). The expected number of nonsynonymous substitutions per gene over 2*N* generations was 0.506, and the ratio of nonsynonymous to synonymous substitutions was 0.0412, which is somewhat less than half the mean value for comparisons of *D. melanogaster* and its close relatives (*e.g*., Campos *et al.* 2017). The major source of these substitutions is the class with 2*Ns_d_* between 2.5 and 4.375, which accounts for 76% of all substitutions, reflecting the fact that mutations in this class have the highest fixation probabilities.

**Figure 3:**
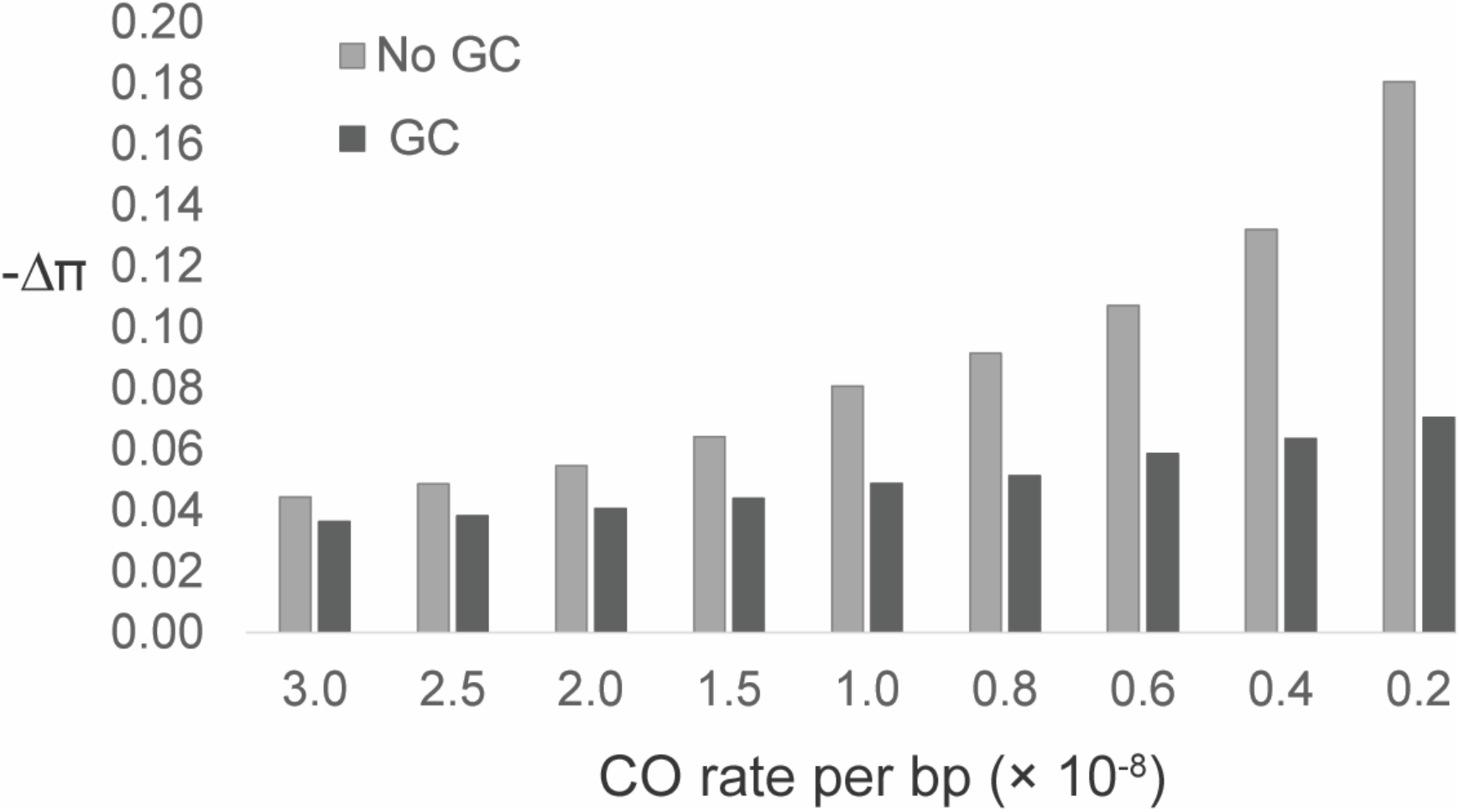
Predicted mean reductions of nucleotide diversity at linked neutral sites compared to neutrality (− Δ*π*), due to recurrent fixations of weakly deleterious semi-dominant mutations with 2.5 ≤ 2*Ns_d_* ≤ 10 in the presence and absence of gene conversion (GC). Results are shown for regions of varying cross-over (CO) rates of recombination. Nucleotide diversity at neutral sites was averaged across a gene comprised of five 300-bp exons and 100-bp introns, in which all intronic sites and 30% of exonic sites were neutral.

The results for single sweeps shown in Table 1 suggest that these theoretical predictions will tend to overestimate sweep effects, so that the results in Figure 3 must be viewed with caution. In addition, the model ignores the mean neutral diversity (*π*) during the progress of a sweep, considering only the contribution from intervals between sweeps. This provides accurate predictions of recurrent sweep effects with 2*Ns_d_* >> 1, since the sweep duration is then a relatively small fraction of the interval between sweeps, unless sweeps are extremely frequent (Charlesworth 2020b). However, this is not true with very weak selection, and the expectation of *π* during a sweep should, therefore, be included in a rigorous treatment. For weak sweeps, it is possible that this quantity could be larger than the expected diversity in the absence of selection, *π*_0_. This follows from the fact that *π* is reduced immediately after the fixation of a neutral mutation at a linked site, but such fixations cannot alter the net expected value of *π*. Since the post-fixation recovery can only bring *π* back up to *π*_0_, the expectation of *π* at linked sites over the course of the neutral fixation must exceed *π*_0_. This arises because, in the absence of recombination, the two alternative haplotypes associated with the two variants in transit to fixation can only coalesce at a time preceding the sweep, and hence have a coalescent time > 2*N.* By continuity, a similar effect must arise with sufficiently weak selection, although it is not seen when 2*Ns_d_* >> 1, when the expected value of *π* is reduced during a sweep (see Figure 8 of Zeng *et al.* 2021), so that the theory used here is likely to overestimate sweep effects. Furthermore, with such weak selection it is somewhat artificial to isolate the effects on *π* of fixations of linked deleterious mutations from the effects of their loss from the population (background selection), given the fact that there will be traffic backwards and forwards between alternative variants at a site that is subject to selection and reverse mutation (Gillespie 1994; Charlesworth & Eyre-Walker 2008).

With the standard *D. melanogaster* sex-averaged rate of crossing over per bp of 1 × 10^−8^, the mean reduction in nucleotide diversity at synonymous sites caused by deleterious sweeps was approximately 5% with gene conversion and 8% in its absence (Figure 3). For the lowest rates of crossing over, and with no gene conversion, average reductions can reach ~19%, suggesting that the fixation of mildly deleterious mutations could play a significant role in organisms or genomic regions with highly reduced rates of recombination. In this low recombination environment however, selective interference will become a factor, and should accelerate the rate of deleterious fixations, while also making deleterious variants behave more like neutral mutations (see below). Regardless of these details, however, it seems unlikely that substitutions of deleterious mutations will have more than a minor effect on average diversity in the *Drosophila* genome overall, particularly compared with the effects of population history, background selection, and sweeps of positively selected mutations. Furthermore, the findings of Mafessoni & Lachmann (2015) suggest that fixations of strongly recessive deleterious mutations will have even smaller effects than those studied here, and can even enhance variability (see Supp. Figure 2, resulting in associative overdominance effects; Charlesworth & Jensen 2021). A detailed study of the possible range of effects of both losses and fixations of weakly selected mutations is planned.

### The effects of interference on deleterious sweeps with *Drosophila-* and human-like parameters

The relatively large input of deleterious mutations could result in interference amongst them, and in turn might affect their probabilities of fixation. In order to evaluate this possibility, genomic regions composed of two adjacent genes with 5 exons and 4 introns each, as well as intergenic regions (see Methods for details) were simulated. Two separate genomic regions were simulated resembling *D. melanogaster* and humans - both in terms of architecture and underlying parameter values - with exons experiencing purifying selection specified by the DFE estimated by Johri *et al.* (2020) for *Drosophila*, and that estimated by Huber *et al.* (2017) for humans. Because interference is more likely to occur in regions of lower recombination rates, additional simulations were carried out for 0.5× and 0.1× the mean recombination rate. Simulations were performed with *N* = 10^4^ and all other parameters were scaled accordingly.

The frequencies of fixation of mildly deleterious mutations were found to be higher than expected with no interference, suggesting an appreciable role of interference (Supp Figure 3). However, the effective reduction in *N*_e_ due to background selection also affects the scaled selection coefficients (Barton 1995; Charlesworth 2012; Campos & Charlesworth 2019). After correcting for the effects of BGS (in which case, expected probabilities were obtained using equations (3) and (4)), the frequencies of fixations of mildly deleterious mutations in humans were found to be unaffected by interference (Supp Figure 4). While no interference effects were observed for regions corresponding to the mean recombination rate in *Drosophila*, for regions with 0.1× the mean recombination rate, the frequencies of fixation were higher than that expected in the absence of interference (Figure 4a).

**Figure 4:**
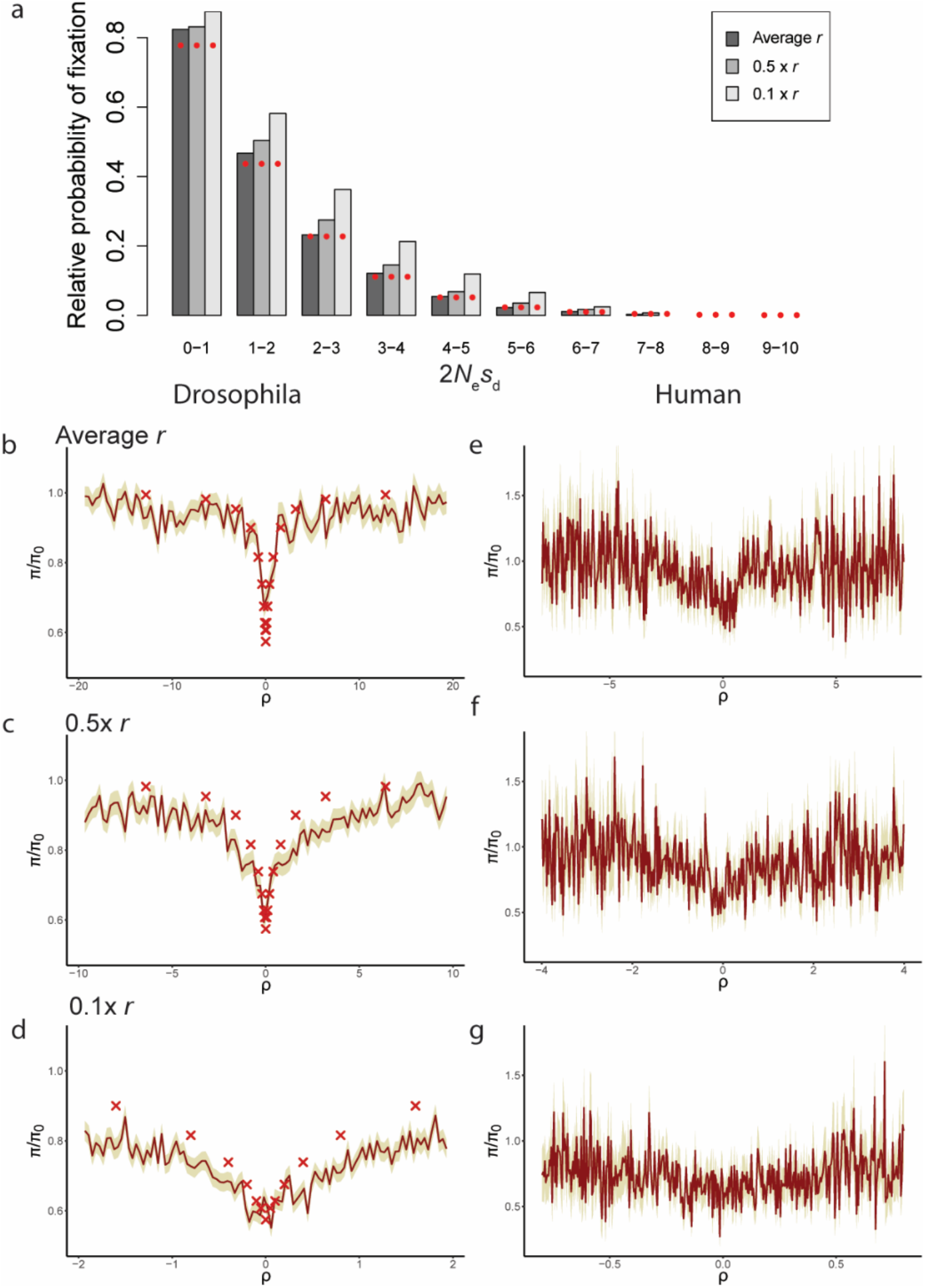
(a) Frequencies of fixation of weakly deleterious mutations relative to that of neutral mutations for different rates of recombination (*r*), when there is a potential for interference amongst deleterious mutations and BGS effects are accounted for. Red solid circles show the expected probability calculated by equation 4 integrating over the interval of 2*N_e_s_d_*. (b-g) Effects of sweeps immediately post-fixation for (b-d) *Drosophila*-like and (e-g) human-like parameters and architecture. Recovery of nucleotide diversity per site (*π*) is shown relative to the mean intergenic diversity under BGS (*π*_0_), around a recent fixation (shown at position 0 on the x-axis). The target site has experienced a weakly deleterious fixation (2*Ns_d_* between 1 and 2). Solid lines represent mean values obtained from all substitutions in all replicates, shaded regions correspond to 1 SE above and 1 SE below the mean. Crosses correspond to simulations based on equation (27) of Tajima (1990) for 2*Ns* = 0. The extent of BGS in these scenarios is: (b) *B* = 0.81, (c) *B* = 0.79, (d) *B* = 0.67, (e) *B* = 0.87, (f) *B* = 0.85, and (g) *B* = 0.83.

Because of these interference effects in regions of low recombination, we further evaluated the times to fixation and the effects of sweeps on linked neutral sites. In both *Drosophila* and humans, the fixation times of deleterious mutations were found to be slightly lengthened in the presence of interference (Supp Figures 5 and 6) as compared to expected time to fixation obtained by numerically integrating Equation 17 of Kimura & Ohta (1969). This suggests that the reduction in *N*_e_ caused by BGS might result in mutations behaving more neutrally, thus taking longer to fix. We have however not accounted for BGS when obtaining expected times numerically. Although the decrease in nucleotide diversity at linked neutral sites remained qualitatively similar to that observed in the absence of interference for *D. melanogaster* (Figure 4b-d, Supp Figure 7), the reduction in mean neutral diversity post-fixation was somewhat more modest for very weakly deleterious alleles in the presence of interference. This smaller effect is probably due to BGS rather than interference per se, as suggested by results above. For recombination rates one-half and one-tenth of the mean, nucleotide diversity up to *ρ*~0.2 was 62% and 60% of the mean intergenic diversity, consistent with a ~40% reduction in variation. For human-like parameters, we similarly observed that the mean nucleotide diversity post-fixation of a weakly deleterious mutation (Figure 4e-g, Supp Figure 7) was very similar to that observed post-fixation of a neutral mutation (Table 1). In sum, for the parameter ranges investigated, we found that interference between weakly deleterious mutations resulted in only very minor deviations from expectations.

### The contribution of deleterious mutations to population- and species-level divergence-based scans

Given the potentially substantial contribution of the weakly deleterious class to observed fixations, it is also of interest to consider their impact on divergence-based analyses (*e.g.*, methodology related to *d_N_*/*d_S_*) and population differentiation-based scans (*e.g.*, methodology related to *F_ST_*). We examined properties related to inferring the proportion of substitutions fixed by positive selection (*α*) by performing MK tests in the presence of mildly deleterious mutations. Specifically, we simulated two scenarios: 1) 50% of mutations at nonsynonymous sites were weakly deleterious, and 50% were neutral, and 2) nonsynonymous sites experienced deleterious mutations that followed the DFE inferred by Johri *et al.* (2020).

Although, as expected, the presence of mildly deleterious mutations substantially increases values of *d_N_*/*d_S_* (Supp Table 7) relative to stronger purifying selection (*i.e*., larger proportions of moderately and strongly deleterious mutations), it also leads to strongly negative values of *α* when performing MK tests. This is due to the fact that mildly deleterious mutations often segregate in the population at low frequency, inflating the total number of segregating nonsynonymous polymorphisms (*P_N_*) significantly (Supp Table 7). As such, the presence of mildly deleterious mutations can result in negative values of *α* in the absence of positive selection. This is consistent with previous studies that have proposed a derived allele frequency cutoff (Fay *et al.* 2001, 2002; Andolfatto 2005; but see Charlesworth & Eyre-Walker 2008) to correct for segregating mildly deleterious alleles, as well as proposed modification of the traditional MK test (*e.g.*, the asymptotic MK test; Messer & Petrov 2013). Nevertheless, under the asymptotic MK test, *α* remains underestimated (Supp Table 7) when the proportion of mildly deleterious mutations is sufficiently high.

In order to study inter-population effects, we simulated a model similar to that recently inferred by Arguello *et al.* (2019), which represents the European and African split of *D. melanogaster* (see Methods), and overlaid it with the estimated DFE of Johri *et al.* (2020). Under this model (*i.e.*, in the absence of positive selection), roughly 50% of SNPs identified as *F_ST_* outliers (defined as representing the upper 1% or 2.5% tails) are mildly deleterious (Figure 5a; Supp Figure 8). Two models of purifying selection are given for comparison (Table 2): 1) a more biologically realistic model in which selection coefficients are scaled to the ancestral population size in defining the DFE classes, such that selection is effectively weaker in the smaller derived population; and 2) an arbitrary model in which the DFE is rescaled such that selective effects are equally strong in the larger ancestral and the smaller derived populations (which, under the chosen demographic model, differ from one another by roughly an order of magnitude). It should be noted that in the model based on Arguello *et al.*, the time post-split between the African and European population is extremely brief, such that there are few or no substitutions post-split (Supp Table 8). Thus, *F_ST_* values are almost entirely dictated by allele frequency differences with respect to co-segregating mutations (Supp Table 8).

**Figure 5:**
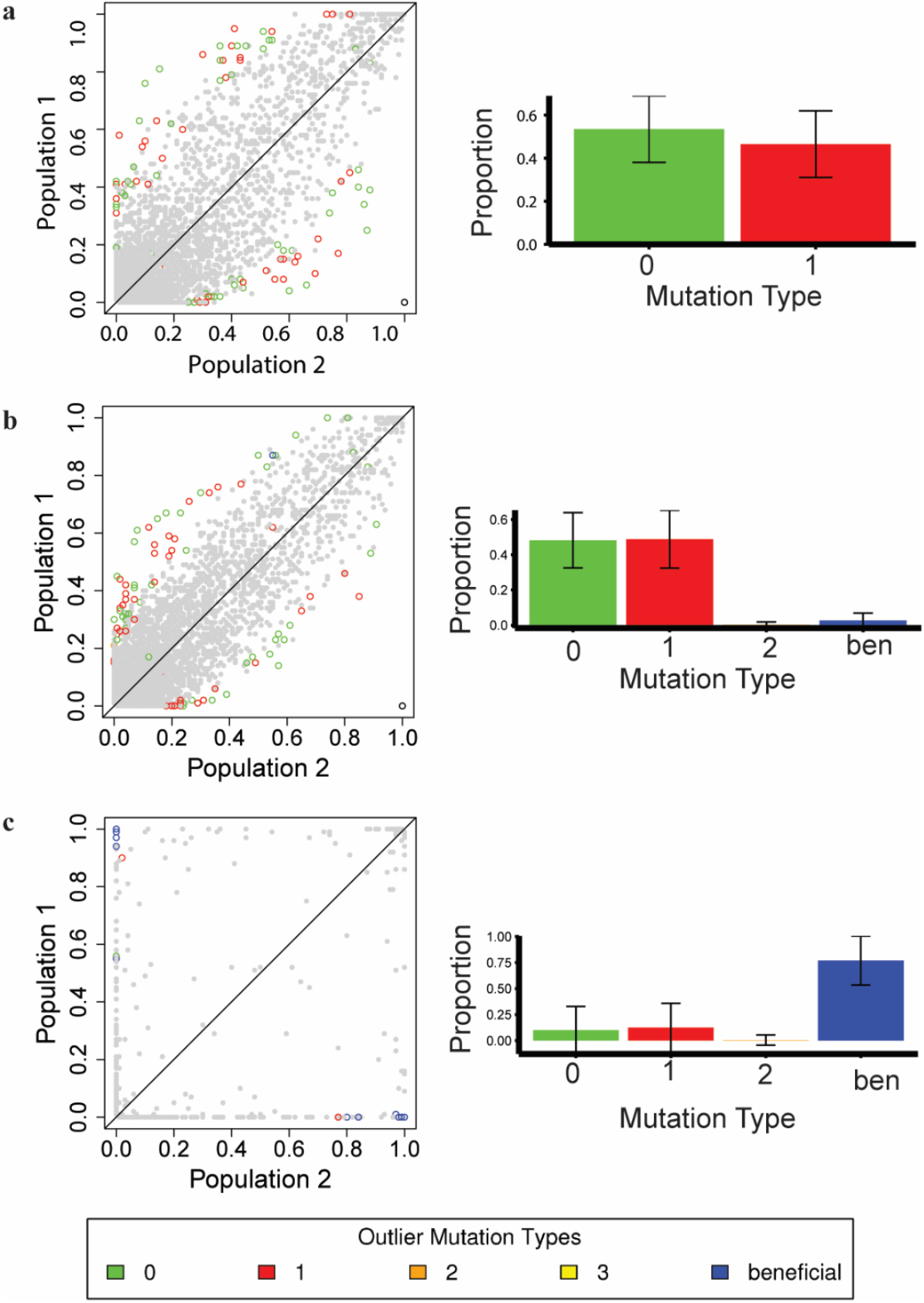
Allele frequencies of SNPs in simulated *D. melanogaster* population 1 (European) *vs* population 2 (African), using parameters of the Arguello *et al.* (2019) model, where the selective effects of all mutations were rescaled with respect to their population sizes after the split (*i.e.*, keeping the strength of selection constant in both populations). Genomic elements experienced (a) purifying selection following the DFE inferred by Johri *et al.* (2020); (b) the same DFE, but with the addition of 1% beneficial mutations with selective effects between 1 < 2*Ns*_*a*_ ≤ 10; or (c) the same DFE, but with the addition of 1% beneficial mutations with selective effects of 2*Ns*_*a*_ = 1000. *Left panel*: Allele frequency plots for 10 (out of 100) replicates simulated. Colored open circles represent *F_ST_* outliers when single SNPs are used to calculate *F_ST_*. Green depicts effectively neutral mutations (belonging to class 0), blue depicts beneficial mutations, and warm colors depict deleterious mutations (belonging to classes 1, 2, and 3), with red representing weakly deleterious mutations. *Right panel*: The distribution of fitness effects of outlier mutations for the corresponding scenarios, showing the mean and standard deviation for all 100 replicates. Sites that were fixed in both populations for the same allele were not included in this analysis.

Mean *F_ST_* values at both neutral and weakly deleterious sites are larger with the rescaled purifying selection model, whereas the unscaled model yields similar values to the purely neutral model (Table 1 and Supp Tables 9, 12). In addition, the frequency of private SNPs at all sites is higher for the rescaled model than the unscaled model, with the neutral model having the lowest value of the three (Supp Table 8). In order to determine whether these weakly deleterious mutations were associated with a decrease in diversity at linked sites, potentially leading to an increase in *F_ST_* values (Charlesworth 1998; Cruickshank & Hahn 2014), we evaluated the relationship between *F_ST_* and nucleotide diversity using neutral variants alone (Supp Figure 9; see also the comparison with directly selected sites in Supp Figures 10-11). The lack of a strong negative correlation suggests that sweep-like effects of deleterious mutations on diversity at linked sites are not primarily responsible for the observed effects on *F_ST_* and the frequency of private alleles. In contrast, there is a negative relationship between nucleotide diversity and *F_ST_* after the strong population bottleneck represented by the neutral model of Li & Stephan (2006).

The lack of sweep-like effects of deleterious mutations is not surprising in view of the time-scale to fixation required for weakly selected mutations, which is of the order of the coalescent time, 2*N* generations; the split times for both the African and European populations are both only a small fraction of their respective coalescent times. Thus, neither new neutral nor weakly deleterious mutations are likely to have reached fixation in either population, consistent with the results shown in the last column of Supp Table 8. In addition, there is no time for large changes in *π* as a result of the altered *N* values.

Instead, it is more likely that BGS effects on the frequencies of segregating mutations explain these patterns (*B* = 0.18 and 0.15 in the African and European populations, respectively, obtained from the ratios of neutral *π* values in the absence of purifying selection to those without, see Supp Table 9). In the absence of rescaling of the DFEs for deleterious mutations (the more biologically plausible case), the enhanced *N* for the African population means that fewer deleterious mutations behave as effectively neutral, so that there is less effect of drift on their frequencies compared with the ancestral population; the reverse is true for the European population. As far as linked neutral variants are concerned, there is likely to be a greater BGS effect in the African than in the ancestral population, and vice-versa for the European population. *F_ST_* relative to the purely neutral case is thus subject to two opposing factors, which presumably explains the lack of any strong effect of this model of purifying selection on *F_ST_* in Table 2 and Supp Table 9. However, the lower effective population size induced by BGS means that rare variants are more likely to be lost after the population split than under neutrality, explaining the increased proportion of private variants compared with neutrality with purifying selection (Supp Table 8). Because of the effects of BGS, the relative *N_e_* values of the African and European populations are less disparate than the relative *N* values, so that the African/European ratio of the proportions of private SNPs is smaller than under neutrality (Supp Table 8).

The effect of rescaling is to keep the proportion of deleterious mutations that are effectively neutral the same in the two descendant populations, with the absolute strength of selection being higher in the African population, so that overall there is a stronger BGS effect in this population. This results in a higher overall fraction of private SNPs, and a larger enrichment of private SNPs in the African population, compared with the neutral case (Supp Table 8). There is a corresponding increase in mean *F_ST_* for both neutral and weakly deleterious variants compared with the neutral case (Table 2 and Supp Table 9).

It is also of interest to consider the contribution of deleterious mutations to outliers (*i.e*., variants that are highly differentiated between the two populations) present in the tails of the distribution in the presence of positive selection. When a class of weakly positively selected sites was added to the DFE (*i.e*. beneficial mutations have fitness effects 1 < 2*Ns*_*a*_ ≤ 10, and comprise 1% of new mutations), these adaptive mutations contributed little to the observed outliers (< 5%) - with mildly deleterious and neutral mutations strongly represented amongst outliers (Figure 5b). Conversely, when positive selection is very strong (*i.e*., mutations have selective effects with 2*Ns*_*a*_=1000, and comprise 1% of new mutations), the majority (~75%) of outliers are drawn from this beneficial class (Figure 5c). Yet, even in the presence of this exceptionally strong and frequent positive selection, ~10% of outliers remain in the mildly deleterious class. Upon simulating the same scenarios - but drawing from the extreme CIs of the demographic parameters to account for underlying uncertainty in the model of Arguello *et al.* (2019) - a very similar proportion of mildly deleterious mutations persists in the presence of weak positive selection (Supp Table 10). However, in the case of strong positive selection, the proportion of strongly beneficial outlier mutations ranges from 25% to 90% depending on the underlying population history (Supp Table 10). It is also noteworthy that under this model of strong, recurrent positive selection, little variation within populations is observed, owing to the severity of the selective sweeps (Supp Table 11), and inter-population allele frequencies are only weakly correlated with one another (Supp Table 8). A haplotype-based measure of population differentiation, *φ_ST_* (Excoffier *et al.* 1992), was also used to identify outliers (*i.e.*, genomic regions present in the tails of the distribution) and was not found to differ substantially (Supp Table 12).

Because any model, including equilibrium neutrality, will have outliers based on empirical *p*-values, we further quantified the properties of outliers (2.5% and 1%) relative to the genome-wide mean. Under models of purifying selection, the values of *F_ST_* outliers were ~2.3-4.5 fold larger than mean *F_ST_* values (Table 2; Supp Table 13), but differed only slightly from the neutral case. Under the strongly bottlenecked neutral model of Li & Stephan (2006), in which the European population experiences a substantial decrease in population size during the bottleneck, the genome-wide mean *F_ST_* values obtained are much higher, as would be expected, though outlier *F_ST_* values were smaller in relative magnitude (~1.5-2.7 fold higher than the means). Under models including weak positive selection, only slight increases in genome-wide *F_ST_* values were observed; whereas the strong positive selection model greatly increased genome-wide values. However, outlier values under both positive selection models were ~2.4-4.6 fold higher than the respective means. Similar results were obtained when using haplotype-based calculations of population differentiation (Supp Table 14). This suggests that recurrent positive selection does not generate substantially larger effect sizes for outlier *F_ST_* values than neutrality or purifying selection.

## CONCLUSIONS

In this paper, we have examined the expected impact of deleterious fixations on polymorphism- and divergence-based scans for selection. Amongst the class of weakly deleterious mutations that have some chance of reaching fixation (1 < 2*Ns_d_* ≤ 10), the resulting sweep effects are highly localized, as expected: on the order of a few dozen base pairs for the parameters considered here. This suggests that deleterious sweeps of this kind are unlikely to be detected in genomic scans based on localized deficits of variation or strongly skewed site frequency spectra. Given the theoretically expected symmetry between beneficial and deleterious sweeps (Maruyama & Kimura 1974; Mafessoni & Lachmann 2015; Charlesworth 2020a), this is expected, as common polymorphism-based methods generally have little power unless selection is exceptionally strong (*e.g.*, Kim & Stephan 2002; Jensen *et al.* 2005; Crisci *et al.* 2013). However, our results suggest that studies that estimate the frequency and strength of classic selective sweeps using patterns of diversity around substitutions (Hernandez *et al.* 2011; Sattath *et al.* 2011; Elyashiv *et al.* 2016), which assume that reductions are entirely caused by the fixation of positively selected mutations, should take into account the effects of reductions caused by neutral and weakly deleterious substitutions.

Furthermore, for among-population comparisons based on *F_ST_*, mildly deleterious mutations contribute significantly to observed outliers, even in the presence of positive selection, particularly in the case of a recent population size reduction. This appears to be the true regardless of whether selective effects are equally strong in both populations (Figure 5), or if selection is relaxed in the smaller derived population (Supp Figure 8). These observations further stress the important point that genomic outliers of a given statistical distribution do not necessarily represent positively selected loci. As such, the performance of outlier-based tests must be assessed on a case-by-case basis under an appropriate baseline model incorporating population history as well as direct and linked purifying selection effects, in order to determine the power and false-positive rates associated with the detection of beneficial alleles (Jensen *et al.* 2019). Moreover, because such deleterious effects are localized to functional sites (*i.e.*, those genomic regions experiencing purifying selection), this may be particularly pernicious in the sense that this class of outlier will not fall in non-functional regions, where they are often attributed to demographic effects. Rather, owing to the common tendency of constructing biological narratives (true or otherwise) around functional outliers (Pavlidis *et al.* 2012), these results suggest that adaptive story-telling may arise from weakly deleterious outliers.

## ACKNOWLEDGMENTS

The authors thank Laurent Excoffier for addressing queries about the AMOVA analyses, as well as Susanne Pfeifer and two anonymous reviewers for providing helpful comments on the manuscript. This work was funded by National Institutes of Health grant R01GM135899 and R35GM139383 to JDJ.

## APPENDIX 1 Calculating the probability of fixation with uniformly distributed fitness effects

For a selection coefficient *s* << 1, the fixation probability of a deleterious mutation when *N_e_* ≠ *N* is approximated by:

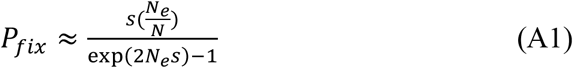

For large *N_e_*, this should be a good approximation, since when 2*N_e_s* ≥ 10, the fixation probability is negligible, so that the contribution from the bin with 2*N_e_s* ≥ 10 can be ignored. For 2*N_e_s* < 10, *s* is sufficiently small that this equation is an accurate approximation.

The integral of this expression over a given interval of 2*N_e_s* values can be found as follows; for convenience, *x* is substituted for *s* and *a* for 2*N_e_*. For *a* > 0, we need to evaluate the following indefinite integral:

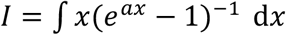

Integration by parts give:

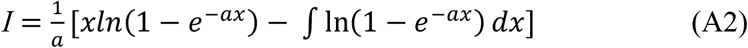

For *a* > 0, the logarithm can be expanded as a power series in exp(−*ax*), which can be integrated term by term:

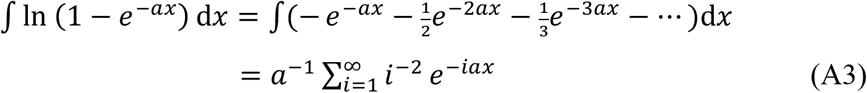

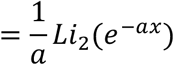

where *Li* is the polylogarithm. The final expression for the integral of equation (1) (provided in the Methods) is thus:

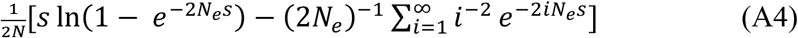

The contribution to the mean fixation probability from the interval *s_i_* to s_*i*+1_ is obtained by dividing this expression by (s_*i*+1_− *s_i_*).

For *s*_0_ = 0, equation (A4) is invalid. However, the integral between 0 and a small positive value of *s*, *s_ε_*, can be found as follows. For 2*N_e_s* << 1, the initial integrand can be approximated by *a*^−1^(1−*ax*/2), so that equation 2 (see Methods) can be replaced with:

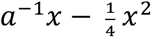

and the indefinite integral of the fixation probability becomes:

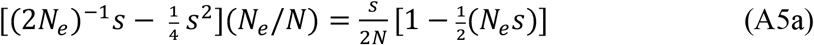

The contribution to the integral of *P_fix_* between *s*_0_ = 0 and *s*_1_ from the interval (0, *s_ε_*) is thus:

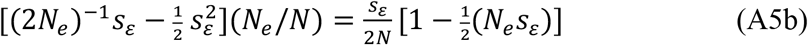

The corresponding mean fixation probability over this interval is:

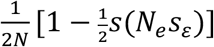

This is slightly smaller than the neutral value, 1/(2*N*), as would be expected when 2*N_e_s* << 1.

Thus, for the interval (*s*_0_, *s*_1_) with *s*_0_ = 0, equation (A5b) should be used for the interval (0≤ *s* ≤ *s_ε_*), and equation (A4) with integration limits *s_ε_* and *s*_1_ for the remainder of the interval.

## SUPPLEMENTARY MATERIAL

**Supp Table 1:** The total number of substitutions with different selective effects observed during simulations that model potential interference amongst deleterious mutations. The simulations were run for 90*N* generations (where *N* = 10^4^ in all cases), with 100 replicates for *D. melanogaster* and 600 replicates for humans.

| Number of substitutions |  |  |  |  |  |  |
| --- | --- | --- | --- | --- | --- | --- |
| *D. melanogaster* |  |  |  | Humans |  |  |
| $2Ns_d$ | Average $r$ | $0.5 \times r$ | $0.1 \times r$ | Average $r$ | $0.5 \times r$ | $0.1 \times r$ |
| (0,1] exon | 31858 | 32772 | 34600 | 7946 | 7932 | 8217 |
| (0,1] inter | 529353 | 534301 | 562314 | 314088 | 315963 | 321766 |
| (1,2] | 4323 | 4684 | 5563 | 133 | 145 | 154 |
| (2,3] | 2554 | 2950 | 4223 | 69 | 86 | 98 |
| (3,4] | 1433 | 1821 | 3027 | 33 | 55 | 60 |
| (4,5] | 825 | 1065 | 2174 | 17 | 23 | 30 |
| (5,6] | 423 | 600 | 1529 | 11 | 10 | 17 |
| (6,7] | 209 | 360 | 1004 | 6 | 5 | 8 |
| (7,8] | 120 | 196 | 692 | 2 | 5 | 4 |
| (8,9] | 56 | 106 | 453 | 2 | 0 | 5 |
| (9,10] | 15 | 61 | 282 | 0 | 0 | 1 |
| $2NBS_d$ | Average $r$ | $0.5 \times r$ | $0.1 \times r$ | Average $r$ | $0.5 \times r$ | $0.1 \times r$ |
| (0,1] inter | 31858 | 32772 | 34600 | 7946 | 7932 | 8217 |
| (0,1] exon | 529353 | 534301 | 562314 | 314088 | 315963 | 321766 |
| (1,2] | 4503 | 4975 | 6809 | 128 | 149 | 154 |
| (2,3] | 2234 | 2713 | 4244 | 58 | 84 | 99 |
| (3,4] | 1168 | 1436 | 2493 | 33 | 39 | 44 |
| (4,5] | 523 | 672 | 1393 | 14 | 12 | 24 |
| (5,6] | 214 | 353 | 768 | 7 | 7 | 9 |
| (6,7] | 104 | 174 | 287 | 2 | 5 | 5 |
| (7,8] | 27 | 70 | 0 | 2 | 0 | 2 |
| (8,9] | 0 | 0 | 0 | 0 | 0 | 1 |
| (9,10] | 0 | 0 | 0 | 0 | 0 | 0 |

**Supp Table 2:**
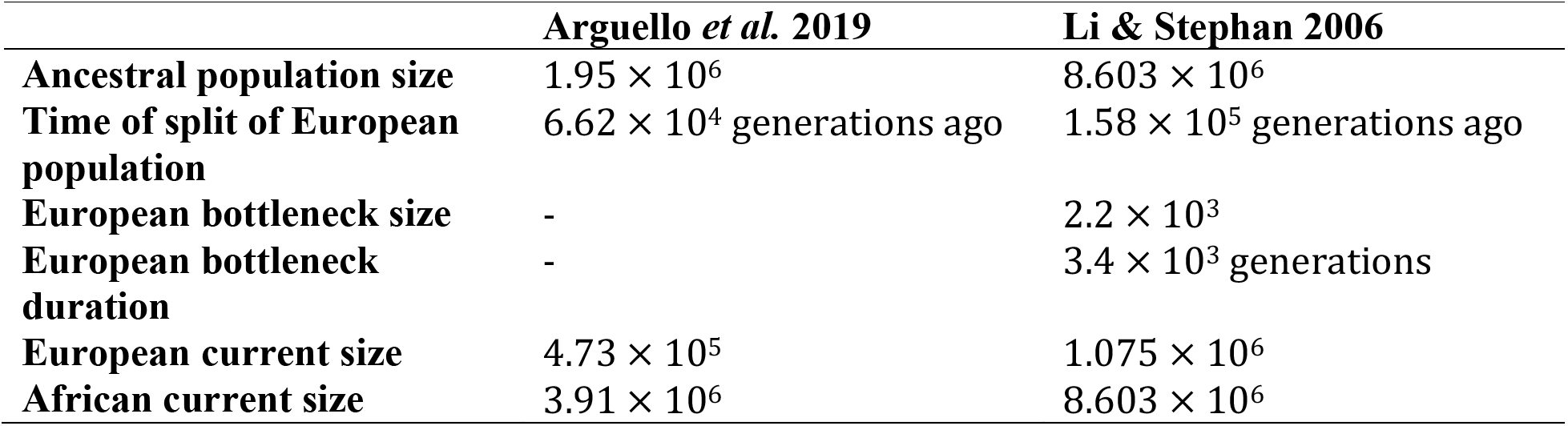
Parameters of the demographic models of *D. melanogaster* simulated for the *F_ST_* analyses.

**Supp Table 3:** The means and standard deviations of fixation times (conditional on fixation) of mutations with different selective effects, obtained from 100 simulated replicates. Fixation times are measured as the time taken for the mutant allele to spread from frequency 1/(2*N*) to frequency 1. Unless otherwise indicated, *h*=0.5.

| Time to fixation (in $2N$ generations) | | | | | | | |
| --- | --- | --- | --- | --- | --- | --- | --- |
| $2Ns$ | min | First quartile (25%) | Median (50%) | Mean | Max | SD | Analytical Mean |
| 0 | 0.57 | 1.30 | 1.82 | 2.08 | 5.77 | 1.04 | 2.00 |
| -1 | 0.45 | 1.20 | 1.67 | 1.96 | 6.19 | 1.05 | 1.97 |
| 1 | 0.58 | 1.14 | 1.52 | 1.73 | 4.09 | 0.99 | 1.97 |
| -5 | 0.53 | 1.14 | 1.36 | 1.55 | 3.37 | 0.74 | 1.56 |
| 5 | 0.58 | 1.15 | 1.49 | 1.66 | 5.42 | 0.57 | 1.56 |
| 5 ($h=0$) | 0.43 | 0.99 | 1.19 | 1.31 | 2.82 | 0.47 | - |
| 5 ($h=1$) | 0.49 | 1.21 | 1.57 | 1.83 | 6.91 | 0.99 | - |
| 10 | 0.50 | 0.77 | 0.97 | 1.04 | 2.18 | 0.36 | 1.11 |
| 20 | 0.33 | 0.56 | 0.70 | 0.73 | 1.40 | 0.23 | 0.70 |
| 30 | 0.25 | 0.44 | 0.51 | 0.53 | 1.05 | 0.14 | 0.53 |
| 40 | 0.23 | 0.38 | 0.43 | 0.44 | 0.80 | 0.10 | 0.42 |

**Supp Table 4:**
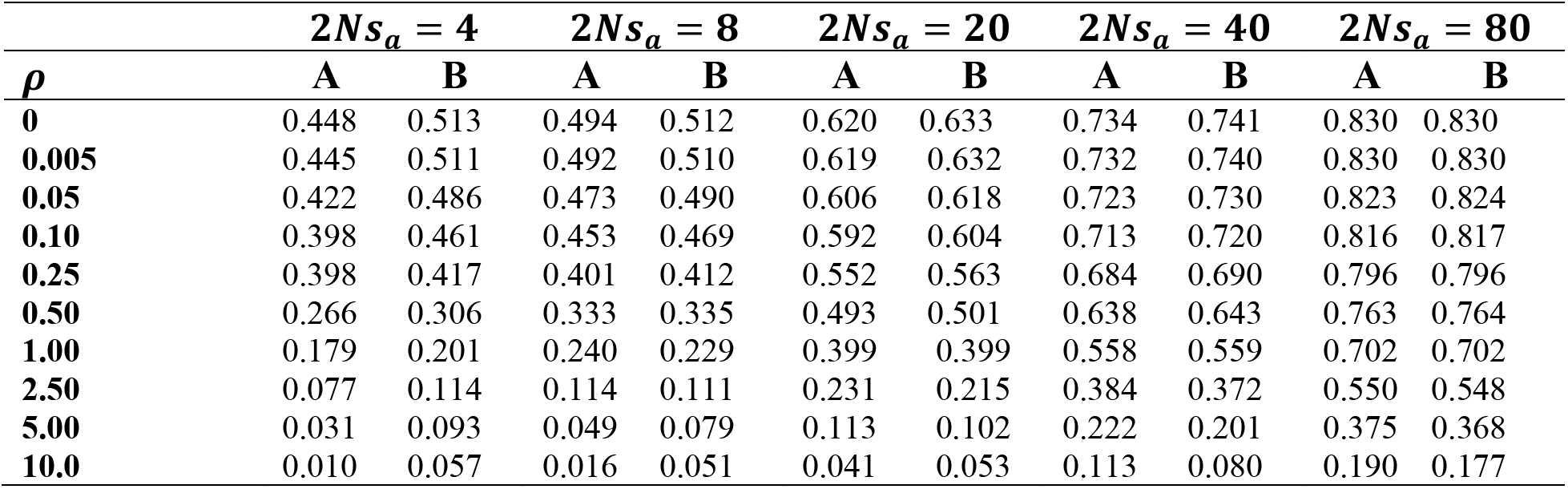
Reduction in diversity (relative to a value of 1 for neutrality) as (A) obtained by Tajima’s (1990) simulations and (B) as approximated by numerical results using equation (14) in Charlesworth 2020b.

**Supp Table 5:** Reduction of diversity (Δ*π*) at *ρ* (= 4*Nr*) distance from the selected site for varying dominance (*h*) and selection coefficients (2*Ns*) calculated using the algorithm of Tajima (1990). For the purpose of these calculations *N* was assumed to be 100 and *θ* = 0.001.

| $\Delta\pi$ | $\rho$ | | | | | | | | | |
| --- | --- | --- | --- | --- | --- | --- | --- | --- | --- | --- |
|  | 0 | 0.05 | 0.1 | 0.2 | 0.4 | 0.8 | 1.6 | 3.2 | 6.4 | 12.8 |
| **$2Ns = 0$; 100000 replicates** |  |  |  |  |  |  |  |  |  |  |
| Mean | 0.4215 | 0.3907 | 0.3635 | 0.3178 | 0.2508 | 0.1706 | 0.0966 | 0.0451 | 0.0178 | 0.0062 |
| SE | 3.3E-04 | 3.5E-04 | 3.6E-04 | 3.8E-04 | 3.9E-04 | 3.6E-04 | 2.7E-04 | 1.6E-04 | 7.1E-05 | 2.4E-05 |
| **$2Ns_a = 2$ [$2Ns_d = 2$]; 100000 replicates** |  |  |  |  |  |  |  |  |  |  |
| $h=0$ Mean | 0.4430 | 0.4157 | 0.3909 | 0.3482 | 0.2825 | 0.1990 | 0.1164 | 0.0556 | 0.0220 | 0.0076 |
| [$h=1$] SE | 3.3E-04 | 3.4E-04 | 3.5E-04 | 3.7E-04 | 3.8E-04 | 3.5E-04 | 2.7E-04 | 1.6E-04 | 7.6E-05 | 3.1E-05 |
| $h=0.1$ Mean | 0.4400 | 0.4121 | 0.3871 | 0.3440 | 0.2781 | 0.1949 | 0.1149 | 0.0547 | 0.0217 | 0.0075 |
| [$h=0.9$] SE | 3.3E-04 | 3.4E-04 | 3.6E-04 | 3.7E-04 | 3.8E-04 | 3.5E-04 | 2.7E-04 | 1.6E-04 | 7.4E-05 | 3.0E-05 |
| $h=0.2$ Mean | 0.4371 | 0.4089 | 0.3836 | 0.3401 | 0.2741 | 0.1913 | 0.1109 | 0.0527 | 0.0208 | 0.0072 |
| [$h=0.8$] SE | 3.3E-04 | 3.5E-04 | 3.6E-04 | 3.7E-04 | 3.8E-04 | 3.5E-04 | 2.7E-04 | 1.6E-04 | 7.4E-05 | 2.7E-05 |
| $h=0.3$ Mean | 0.4344 | 0.4057 | 0.3801 | 0.3364 | 0.2704 | 0.1880 | 0.1085 | 0.0514 | 0.0203 | 0.0070 |
| [$h=0.7$] SE | 3.3E-04 | 3.5E-04 | 3.6E-04 | 3.8E-04 | 3.8E-04 | 3.5E-04 | 2.7E-04 | 1.6E-04 | 7.4E-05 | 2.9E-05 |
| $h=0.4$ Mean | 0.4317 | 0.4029 | 0.3769 | 0.3328 | 0.2664 | 0.1843 | 0.1061 | 0.0501 | 0.0198 | 0.0069 |
| [$h=0.6$] SE | 3.3E-04 | 3.5E-04 | 3.6E-04 | 3.8E-04 | 3.9E-04 | 3.5E-04 | 2.7E-04 | 1.6E-04 | 7.3E-05 | 2.9E-05 |
| $h=0.5$ Mean | 0.4289 | 0.3996 | 0.3734 | 0.3290 | 0.2623 | 0.1807 | 0.1035 | 0.0488 | 0.0192 | 0.0067 |
| [$h=0.5$] SE | 3.3E-04 | 3.5E-04 | 3.6E-04 | 3.8E-04 | 3.9E-04 | 3.5E-04 | 2.7E-04 | 1.6E-04 | 7.2E-05 | 2.6E-05 |
| $h=0.6$ Mean | 0.4264 | 0.3967 | 0.3699 | 0.3251 | 0.2585 | 0.1773 | 0.1011 | 0.0474 | 0.0187 | 0.0065 |
| [$h=0.4$] SE | 3.3E-04 | 3.5E-04 | 3.6E-04 | 3.8E-04 | 3.9E-04 | 3.5E-04 | 2.7E-04 | 1.6E-04 | 7.2E-05 | 2.8E+00 |
| $h=0.7$ Mean | 0.4236 | 0.3935 | 0.3665 | 0.3210 | 0.2545 | 0.1737 | 0.0987 | 0.0461 | 0.0182 | 0.0064 |
| [$h=0.3$] SE | 3.3E-04 | 3.5E-04 | 3.6E-04 | 3.8E-04 | 3.9E-04 | 3.5E-04 | 2.6E-04 | 1.5E-04 | 7.1E-05 | 2.8E-05 |
| $h=0.8$ Mean | 0.4211 | 0.4032 | 0.3769 | 0.3326 | 0.2673 | 0.1896 | 0.0963 | 0.0449 | 0.0177 | 0.0062 |
| [$h=0.2$] SE | 3.3E-04 | 3.3E-04 | 3.5E-04 | 3.6E-04 | 3.7E-04 | 3.3E-04 | 2.6E-04 | 1.5E-04 | 7.1E-05 | 3.1E-05 |
| $h=0.9$ Mean | 0.4218 | 0.3868 | 0.3595 | 0.3132 | 0.2463 | 0.1666 | 0.0938 | 0.0436 | 0.0171 | 0.0060 |
| [$h=0.1$] SE | 3.4E-04 | 3.5E-04 | 3.6E-04 | 3.8E-04 | 3.9E-04 | 3.5E-04 | 2.6E-04 | 1.5E-04 | 7.1E-05 | 3.0E-05 |
| $h=1$ Mean | 0.4158 | 0.3842 | 0.3562 | 0.3097 | 0.2429 | 0.1633 | 0.0915 | 0.0425 | 0.0167 | 0.0059 |
| [$h=0$] SE | 3.3E-04 | 3.5E-04 | 3.7E-04 | 3.8E-04 | 3.9E-04 | 3.5E-04 | 2.6E-04 | 1.5E-04 | 7.1E-05 | 2.7E-05 |
| **$2Ns_a = 5$ [$2Ns_d = 5$]; 1000 replicates** |  |  |  |  |  |  |  |  |  |  |
| $h=0$ Mean | 0.4916 | 0.4724 | 0.4422 | 0.4006 | 0.3516 | 0.2554 | 0.1544 | 0.0812 | 0.0321 | 0.0109 |
| [$h=1$] SE | 3.1E-03 | 3.2E-03 | 3.3E-03 | 3.3E-03 | 3.6E-03 | 3.4E-03 | 2.8E-03 | 1.8E-03 | 8.2E-04 | 3.1E-04 |
| $h=0.9$ Mean | 0.4249 | 0.4022 | 0.3755 | 0.3357 | 0.2712 | 0.1899 | 0.1119 | 0.0523 | 0.0213 | 0.0074 |
| [$h=0.1$] SE | 3.5E-03 | 3.7E-03 | 3.7E-03 | 4.0E-03 | 4.1E-03 | 3.7E-03 | 2.7E-03 | 1.7E-03 | 7.9E-04 | 3.0E-04 |
| $h=1$ Mean | 0.4201 | 0.3932 | 0.3717 | 0.3268 | 0.2677 | 0.1774 | 0.1046 | 0.0530 | 0.0185 | 0.0068 |
| [$h=0$] SE | 3.5E-04 | 3.5E-03 | 3.9E-03 | 3.9E-03 | 4.2E-03 | 3.7E-03 | 2.8E-03 | 1.7E-03 | 7.7E-04 | 3.0E-04 |
| **$2Ns_a = 10$ [$2Ns_d = 10$]; 1000 replicates** |  |  |  |  |  |  |  |  |  |  |
| $h=0$ Mean | 0.5701 | 0.4724 | 0.4422 | 0.4006 | 0.3516 | 0.2554 | 0.1544 | 0.0812 | 0.0321 | 0.0109 |
| [$h=1$] SE | 2.5E-03 | 3.2E-03 | 3.3E-03 | 3.3E-03 | 3.6E-03 | 3.4E-03 | 2.8E-03 | 1.8E-03 | 8.2E-04 | 3.1E-04 |
| $h=1$ Mean | 0.4507 | 0.4315 | 0.4124 | 0.3796 | 0.3231 | 0.2452 | 0.1548 | 0.0794 | 0.0319 | 0.0109 |
| [$h=0$] SE | 1.2E-03 | 1.2E-03 | 1.2E-03 | 1.3E-03 | 1.3E-03 | 1.2E-03 | 9.8E-04 | 6.1E-04 | 2.9E-04 | 1.1E-04 |

**Supp Table 6:** Distortion of the SFS at neutral sites linked to the selected site at a distance measured by *ρ*, and calculated immediately after the fixation of a weakly selected mutation. 100 replicates were used to obtain these values. Tajima’s *D* was calculated in non-overlapping sliding windows of 50 base pairs, with the distance from the selected site corresponding to the mid-point of the window.

| $\rho$ | Number of bases from the selected site in *Drosophila* | $2Ns = 0$ | | $2Ns = +5$ | | $2Ns = -5$ | |
| --- | --- | --- | --- | --- | --- | --- | --- |
| | | Tajima's $D$<br>mean | Tajima's $D$<br>SE | Tajima's $D$<br>mean | Tajima's $D$<br>SE | Tajima's $D$<br>mean | Tajima's $D$<br>SE |
| 1 | 25 | -0.1493 | 0.0712 | -0.2097 | 0.0671 | -0.1852 | 0.0867 |
| 3 | 75 | -0.0979 | 0.0696 | -0.0827 | 0.0717 | -0.0476 | 0.0919 |
| 5 | 125 | 0.00165 | 0.0731 | -0.1042 | 0.0712 | -0.0176 | 0.0871 |
| 7 | 175 | -0.0355 | 0.0657 | -0.0731 | 0.0699 | -0.0193 | 0.0869 |
| 9 | 225 | -0.0864 | 0.0712 | -0.0214 | 0.0729 | -0.1359 | 0.0816 |

**Supp Table 7:** Number of polymorphic (*P*) and fixed (*D*) sites at nonsynonymous (*N*) and synonymous (*S*) sites, and the estimated proportion of substitutions fixed by positive selection (*α*), for an increasing proportion of mildly deleterious mutations at nonsynonymous sites. The ratio of the rate of divergence at nonsynonymous relative to that at synonymous sites is denoted by *d*_*N*_/*d*_*S*_.

| | % weakly deleterious NS sites | $P_N$ | $D_N$ | $P_S$ | $D_S$ | $d_N/d_S$ | $\alpha$ | $\alpha_{asym}$<br>[ 95% CI] |
| --- | --- | --- | --- | --- | --- | --- | --- | --- |
| Neutral | 0% | 522 | 439 | 255 | 221 | 0.99 | -0.05 | -0.06 [-0.21, 0.09] |
| Weakly deleterious | 50% | 447 | 261 | 240 | 237 | 0.55 | -0.58 | -0.35 [-0.72, 0.03] |
| Johri *et al.*<br>DFE | 66% | 281 | 60 | 173 | 254 | 0.12 | -5.22 | -3.55 [-5.10, -2.01] |

**Supp Table 8:** Composition of shared and private SNPs between populations (calculated in sliding windows of 500 bp).

|  | **Fraction of shared SNPs** | **Fraction of private SNPs in African population** | **Fraction of private SNPs in European population** | **Fraction of fixed differences** |
| --- | --- | --- | --- | --- |
| **Neutral** | 0.538 | 0.355 | 0.108 | 0.000 |
| **Purifying selection (rescaled)** | 0.363 | 0.521 | 0.116 | 0.000 |
| **Purifying selection (not rescaled)** | 0.417 | 0.440 | 0.143 | 0.000 |
| **Weak positive selection** | 0.414 | 0.442 | 0.145 | 0.000 |
| **Strong positive selection** | 0.047 | 0.617 | 0.334 | 0.002 |
| **Neutral bottleneck** | 0.242 | 0.706 | 0.052 | 0.000 |

**Supp Table 9:**
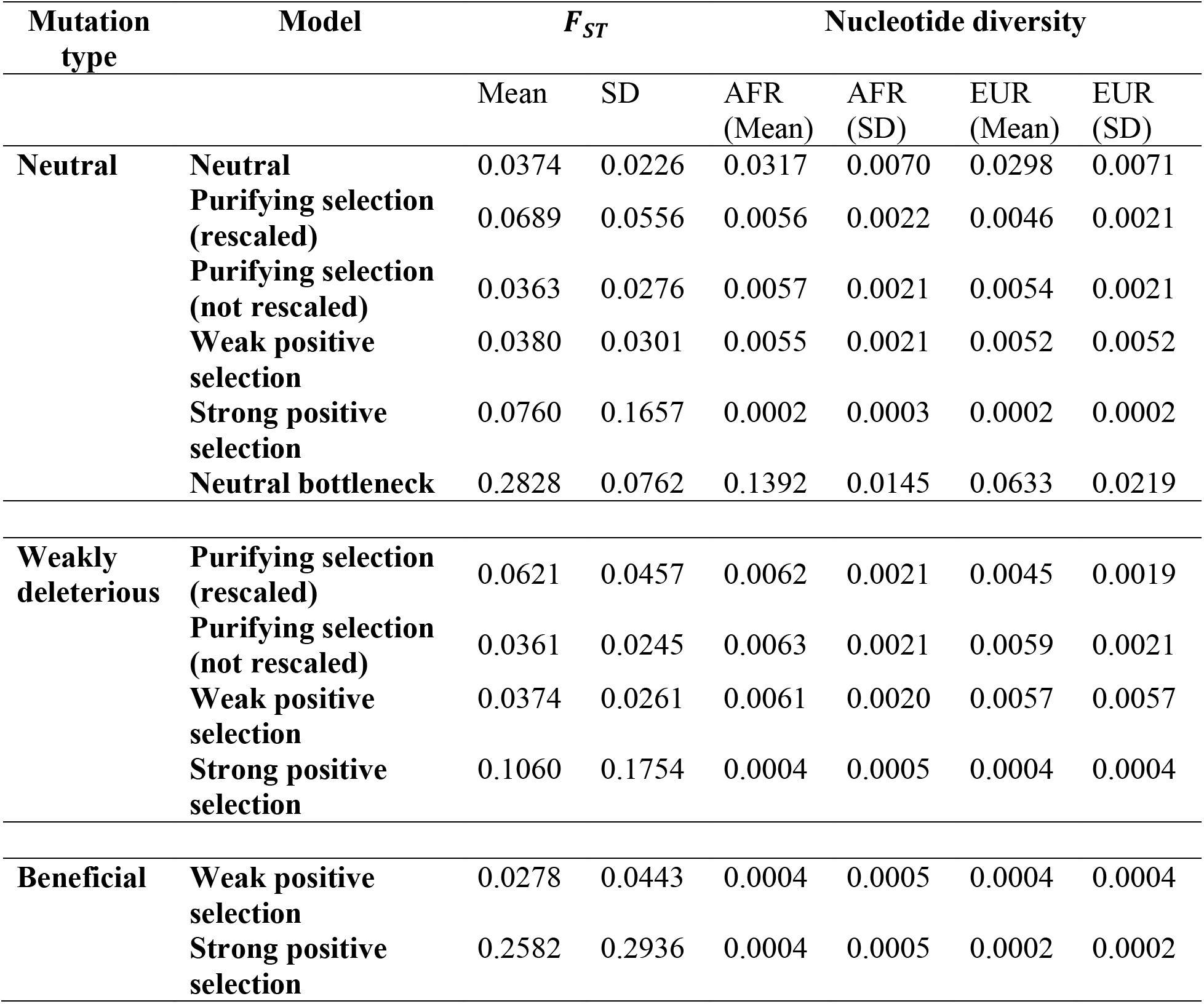
Mean *F_ST_* and nucleotide diversity in the African and European population, for neutral, weakly deleterious, and beneficial mutations separately, calculated for windows of 500 bp.

**Supp Table 10:** The proportion of mutation types amongst the 1% outliers under various evolutionary scenarios, when simulations were performed with the maximum and minimum values belonging to the CIs of estimates of population sizes and the time of split as presented in Arguello *et al.* 2019.

| Model | Time of split | African $N$ | European $N$ | Proportion of mutation types in outliers | | | | |
| --- | --- | --- | --- | --- | --- | --- | --- | --- |
| | | | | $f_0$ | $f_1$ | $f_2$ | $f_3$ | Beneficial |
| Purifying selection | 1.03x10<sup>5</sup> | 3.02x10<sup>6</sup> | 2.03x10<sup>5</sup> | 0.488 | 0.512 | 0.000 | 0.000 | 0.000 |
|  |  | 4.69x10<sup>6</sup> | 3.89x10<sup>6</sup> | 0.531 | 0.462 | 0.008 | 0.000 | 0.000 |
|  |  | 3.02x10<sup>6</sup> | 3.89x10<sup>6</sup> | 0.505 | 0.486 | 0.009 | 0.000 | 0.000 |
|  |  | 4.69x10<sup>6</sup> | 2.03x10<sup>5</sup> | 0.534 | 0.456 | 0.010 | 0.000 | 0.000 |
|  | 1.17x10<sup>4</sup> | 3.02x10<sup>6</sup> | 2.03x10<sup>5</sup> | 0.484 | 0.505 | 0.011 | 0.000 | 0.000 |
|  |  | 4.69x10<sup>6</sup> | 3.89x10<sup>6</sup> | 0.591 | 0.409 | 0.000 | 0.000 | 0.000 |
|  |  | 3.02x10<sup>6</sup> | 3.89x10<sup>6</sup> | 0.470 | 0.523 | 0.008 | 0.000 | 0.000 |
|  |  | 4.69x10<sup>6</sup> | 2.03x10<sup>5</sup> | 0.510 | 0.479 | 0.010 | 0.000 | 0.000 |
| Strong positive selection | 1.03x10<sup>5</sup> | 3.02x10<sup>6</sup> | 2.03x10<sup>5</sup> | 0.000 | 0.118 | 0.000 | 0.000 | 0.882 |
|  |  | 4.69x10<sup>6</sup> | 3.89x10<sup>6</sup> | 0.038 | 0.077 | 0.000 | 0.038 | 0.846 |
|  |  | 3.02x10<sup>6</sup> | 3.89x10<sup>6</sup> | 0.080 | 0.080 | 0.000 | 0.000 | 0.840 |
|  |  | 4.69x10<sup>6</sup> | 2.03x10<sup>5</sup> | 0.050 | 0.050 | 0.000 | 0.000 | 0.900 |
|  | 1.17x10<sup>4</sup> | 3.02x10<sup>6</sup> | 2.03x10<sup>5</sup> | 0.188 | 0.375 | 0.000 | 0.000 | 0.438 |
|  |  | 4.69x10<sup>6</sup> | 3.89x10<sup>6</sup> | 0.286 | 0.381 | 0.000 | 0.000 | 0.333 |
|  |  | 3.02x10<sup>6</sup> | 3.89x10<sup>6</sup> | 0.313 | 0.375 | 0.000 | 0.000 | 0.313 |
|  |  | 4.69x10<sup>6</sup> | 2.03x10<sup>5</sup> | 0.250 | 0.500 | 0.000 | 0.000 | 0.250 |
| Weak Positive selection | 1.03x10<sup>5</sup> | 3.02x10<sup>6</sup> | 2.03x10<sup>5</sup> | 0.505 | 0.462 | 0.000 | 0.000 | 0.032 |
|  |  | 4.69x10<sup>6</sup> | 3.89x10<sup>6</sup> | 0.438 | 0.512 | 0.000 | 0.000 | 0.050 |
|  |  | 3.02x10<sup>6</sup> | 3.89x10<sup>6</sup> | 0.438 | 0.504 | 0.000 | 0.000 | 0.058 |
|  |  | 4.69x10<sup>6</sup> | 2.03x10<sup>5</sup> | 0.436 | 0.543 | 0.011 | 0.000 | 0.011 |
|  | 1.17x10<sup>4</sup> | 3.02x10<sup>6</sup> | 2.03x10<sup>5</sup> | 0.387 | 0.602 | 0.000 | 0.000 | 0.011 |
|  |  | 4.69x10<sup>6</sup> | 3.89x10<sup>6</sup> | 0.431 | 0.549 | 0.020 | 0.000 | 0.000 |
|  |  | 3.02x10<sup>6</sup> | 3.89x10<sup>6</sup> | 0.391 | 0.609 | 0.000 | 0.000 | 0.000 |
|  |  | 4.69x10<sup>6</sup> | 2.03x10<sup>5</sup> | 0.465 | 0.488 | 0.000 | 0.000 | 0.047 |

Results are presented for 10 replicates of each scenario. The DFE was not scaled in any of the above scenarios.

**Supp Table 11:** Average nucleotide diversity (*π*) per site of African and European populations simulated for *F_ST_* analyses.

| Model | Population | $\pi$ | |
| --- | --- | --- | --- |
|  |  | Mean | Standard deviation |
| Neutral | African | 0.0317 | 0.0070 |
|  | European | 0.0298 | 0.0071 |
| Weak positive selection | African | 0.0121 | 0.0033 |
|  | European | 0.0114 | 0.0033 |
| Strong positive selection | African | 0.0011 | 0.0008 |
|  | European | 0.0008 | 0.0007 |
| Purifying selection (DFE rescaled) | African | 0.0118 | 0.0033 |
|  | European | 0.0091 | 0.0031 |
| Purifying selection (DFE not rescaled) | African | 0.0120 | 0.0032 |
|  | European | 0.0113 | 0.0033 |
| Neutral bottleneck | African | 0.1392 | 0.0145 |
|  | European | 0.0633 | 0.0219 |

**Supp Table 12:** Proportions of different classes of selected mutations in the set of all new mutations (referred to as “input”), all segregating mutations, and those observed amongst the 1% outliers detected using the distribution of *φ_ST_*.

| Evolutionary Model | Window size | Proportion of mutations |  |  |  |  |
| --- | --- | --- | --- | --- | --- | --- |
|  |  | Neutral | Weakly deleterious | Moderately deleterious | Strongly deleterious | Beneficial |
| Purifying selection | Input | 0.25 | 0.49 | 0.04 | 0.22 | - |
|  | Segregating | 0.3825 | 0.5918 | 0.0205 | 0.0052 | - |
|  | Outliers (500 bp) | 0.3857 | 0.5899 | 0.0201 | 0.0043 | - |
|  | Outliers (1000bp) | 0.3867 | 0.5946 | 0.0130 | 0.0057 | - |
|  | Outliers (2000bp) | 0.3718 | 0.6054 | 0.0165 | 0.0064 | - |
| Purifying selection (rescaled DFE) | Input | 0.25 | 0.49 | 0.04 | 0.22 | - |
|  | Segregating | 0.3908 | 0.5904 | 0.0175 | 0.0014 | - |
|  | Outliers (500 bp) | 0.3855 | 0.5921 | 0.0205 | 0.0019 | - |
|  | Outliers (1000bp) | 0.3852 | 0.5945 | 0.0191 | 0.0012 | - |
|  | Outliers (2000bp) | 0.3837 | 0.5957 | 0.0198 | 0.0007 | - |
| With 1% weak positive selection | Input | 0.246 | 0.489 | 0.039 | 0.217 | 0.010 |
|  | Segregating | 0.3754 | 0.5777 | 0.0206 | 0.0058 | 0.0205 |
|  | Outliers (500 bp) | 0.3683 | 0.5894 | 0.0170 | 0.0070 | 0.0182 |
|  | Outliers (1000bp) | 0.3742 | 0.5821 | 0.0177 | 0.0077 | 0.0184 |
|  | Outliers (2000bp) | 0.3765 | 0.5757 | 0.0219 | 0.0064 | 0.0196 |
| With 1% strong positive selection | Input | 0.246 | 0.489 | 0.039 | 0.217 | 0.010 |
|  | Segregating | 0.2589 | 0.5444 | 0.0354 | 0.0252 | 0.1361 |
|  | Outliers (500 bp) | 0.2438 | 0.4962 | 0.0325 | 0.0433 | 0.1842 |
|  | Outliers (1000bp) | 0.2642 | 0.4884 | 0.0296 | 0.0361 | 0.1817 |
|  | Outliers (2000bp) | 0.2580 | 0.5135 | 0.0319 | 0.0344 | 0.1622 |

**Supp Table 13:** Mean genome-wide and outlier *F_ST_* values for European and African *D. melanogaster* populations (expanded from Table 2) - for window sizes of both 10 and 20 SNPs, and the 1% and 2.5% tails of the distribution.

| Evolutionary model | Window size | Mean $F_{ST}$ | Outlier $F_{ST}$ (2.5%) | Outlier $F_{ST}$ (1%) | Outlier $F_{ST}$ (2.5%) / Mean $F_{ST}$ | Outlier $F_{ST}$ (1%) / Mean $F_{ST}$ |
| --- | --- | --- | --- | --- | --- | --- |
| Neutral | 10 SNPs | 0.0363 | 0.1585 | 0.1628 | 4.3662 | 4.4838 |
|  | 20 SNPs | 0.0369 | 0.1411 | 0.1471 | 3.8203 | 3.9844 |
|  | 500 bp | 0.0374 | 0.1144 | 0.1336 | 3.0596 | 3.5734 |
| Purifying selection (rescaled DFE) | 10 SNPs | 0.0613 | 0.2497 | 0.2637 | 4.0707 | 4.2999 |
|  | 20 SNPs | 0.0646 | 0.2234 | 0.2234 | 3.4564 | 3.4564 |
|  | 500 bp | 0.0666 | 0.2452 | 0.2982 | 3.6832 | 4.4795 |
| Purifying selection | 10 SNPs | 0.0352 | 0.1467 | 0.1454 | 4.1673 | 4.1296 |
|  | 20 SNPs | 0.0361 | 0.1311 | 0.1311 | 3.6288 | 3.6288 |
|  | 500 bp | 0.0365 | 0.1115 | 0.1290 | 3.0550 | 3.5328 |
| Neutral bottleneck | 10 SNPs | 0.2557 | 0.6459 | 0.6956 | 2.5260 | 2.7204 |
|  | 20 SNPs | 0.2681 | 0.6071 | 0.6560 | 2.2650 | 2.4472 |
|  | 500 bp | 0.2828 | 0.4688 | 0.4922 | 1.6576 | 1.7403 |
| With weak positive selection | 10 SNPs | 0.0362 | 0.1590 | 0.1631 | 4.3932 | 4.5057 |
|  | 20 SNPs | 0.0374 | 0.1355 | 0.1355 | 3.6270 | 3.6270 |
|  | 500 bp | 0.0378 | 0.1228 | 0.1443 | 3.2452 | 3.8143 |
| With strong positive selection | 10 SNPs | 0.2347 | 0.6689 | 0.6689 | 2.8507 | 2.8507 |
|  | 20 SNPs | 0.2954 | 0.6403 | 0.6403 | 2.1677 | 2.1677 |
|  | 500 bp | 0.2353 | 0.8685 | 0.9108 | 3.6908 | 3.8705 |

**Supp Table 14:**
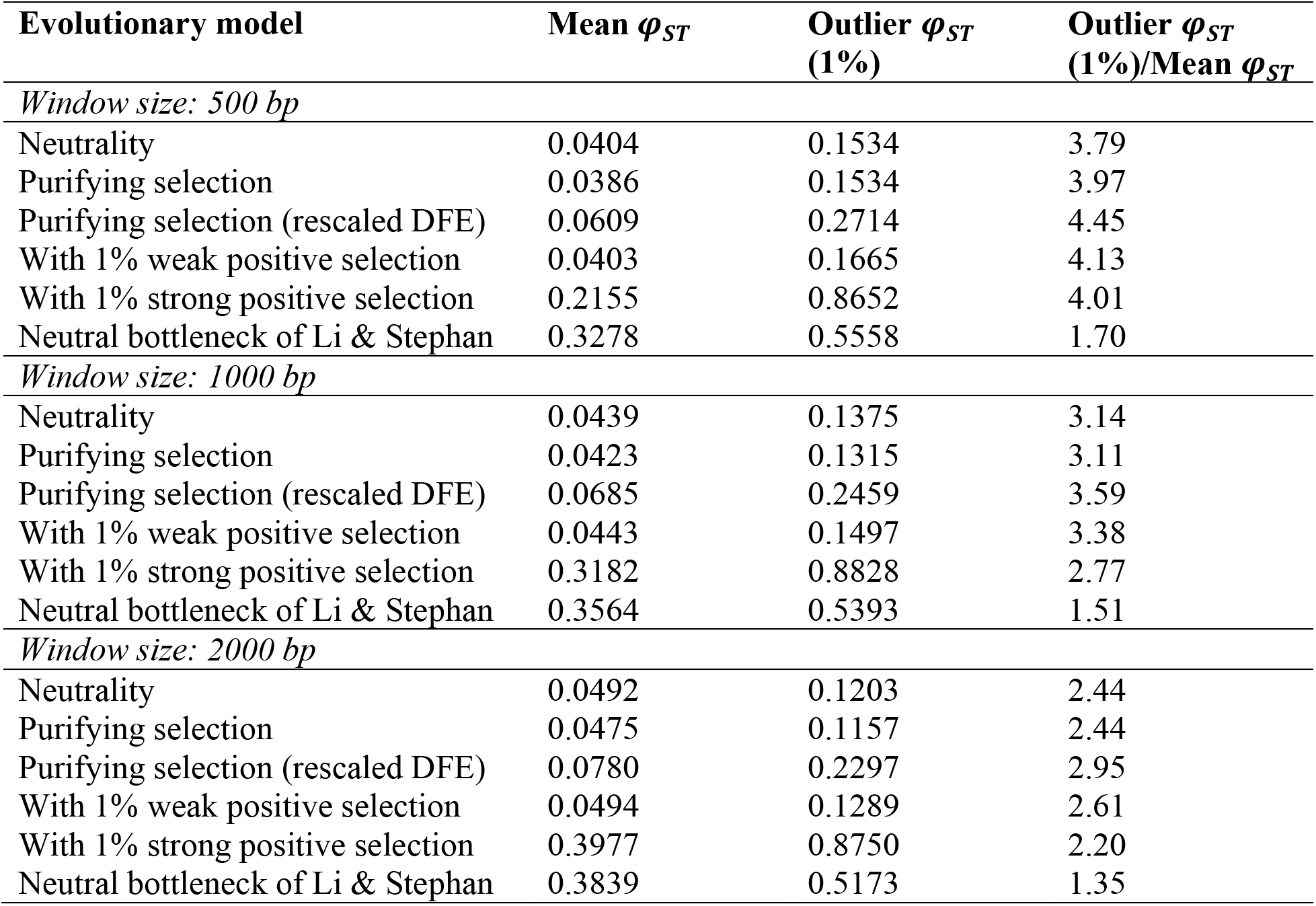
Mean genome-wide and outlier *φ_ST_* values for European and African *D. melanogaster* populations-for constant window sizes of 500, 1000 and 2000 bases, and the 1% tails of the distribution.

| Evolutionary model | Mean $\varphi_{ST}$ | Outlier $\varphi_{ST}$<br>(1%) | Outlier $\varphi_{ST}$<br>(1%)/Mean $\varphi_{ST}$ |
| --- | --- | --- | --- |
| *Window size: 500 bp* |  |  |  |
| Neutrality | 0.0404 | 0.1534 | 3.79 |
| Purifying selection | 0.0386 | 0.1534 | 3.97 |
| Purifying selection (rescaled DFE) | 0.0609 | 0.2714 | 4.45 |
| With 1% weak positive selection | 0.0403 | 0.1665 | 4.13 |
| With 1% strong positive selection | 0.2155 | 0.8652 | 4.01 |
| Neutral bottleneck of Li & Stephan | 0.3278 | 0.5558 | 1.70 |
| *Window size: 1000 bp* |  |  |  |
| Neutrality | 0.0439 | 0.1375 | 3.14 |
| Purifying selection | 0.0423 | 0.1315 | 3.11 |
| Purifying selection (rescaled DFE) | 0.0685 | 0.2459 | 3.59 |
| With 1% weak positive selection | 0.0443 | 0.1497 | 3.38 |
| With 1% strong positive selection | 0.3182 | 0.8828 | 2.77 |
| Neutral bottleneck of Li & Stephan | 0.3564 | 0.5393 | 1.51 |
| *Window size: 2000 bp* |  |  |  |
| Neutrality | 0.0492 | 0.1203 | 2.44 |
| Purifying selection | 0.0475 | 0.1157 | 2.44 |
| Purifying selection (rescaled DFE) | 0.0780 | 0.2297 | 2.95 |
| With 1% weak positive selection | 0.0494 | 0.1289 | 2.61 |
| With 1% strong positive selection | 0.3977 | 0.8750 | 2.20 |
| Neutral bottleneck of Li & Stephan | 0.3839 | 0.5173 | 1.35 |

**Supp Figure 1:**
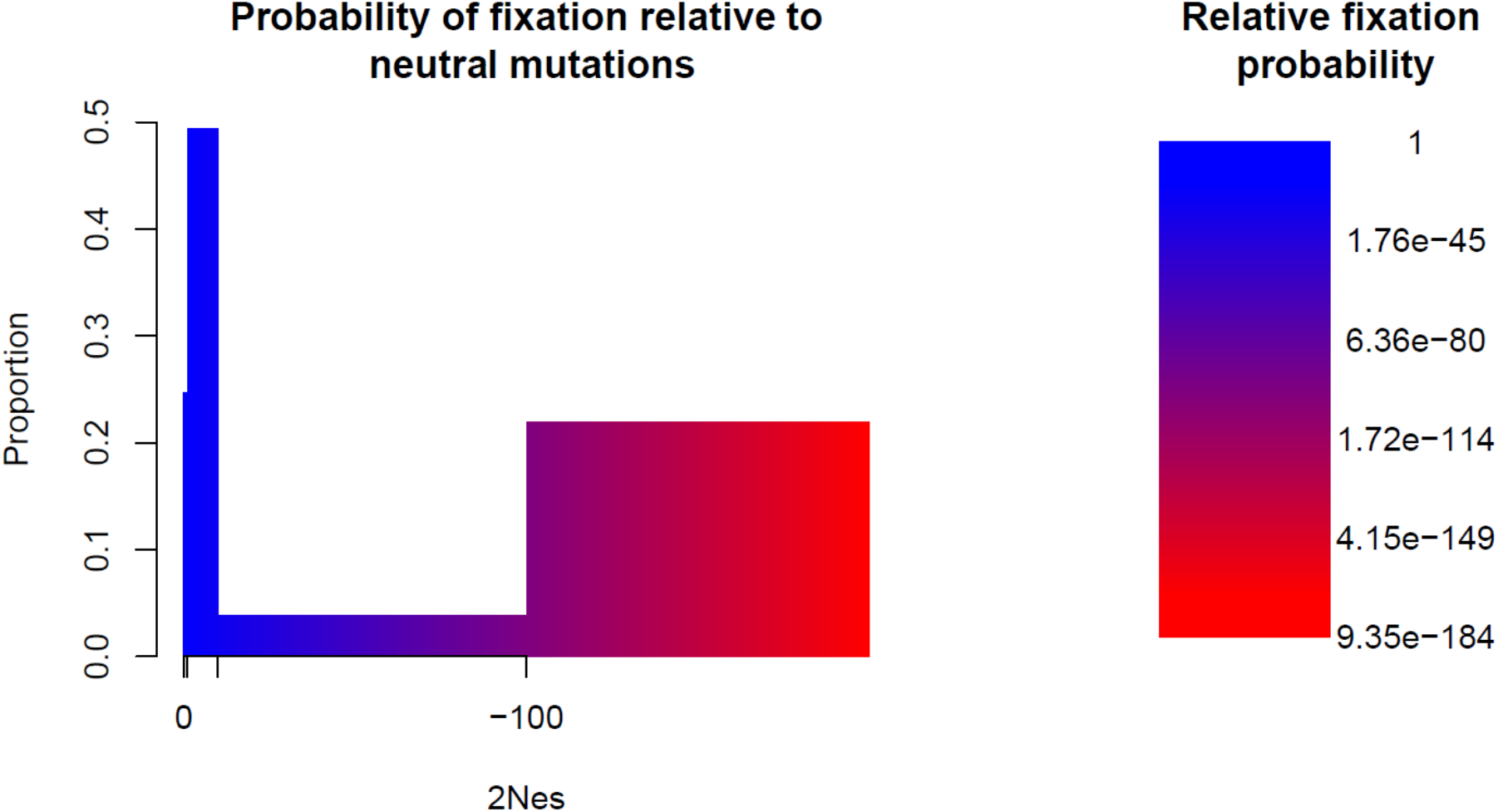
The DFE of deleterious mutations assumed in this study (from Johri *et al.* 2020), color-coded to display the probabilities of fixation of deleterious mutations relative to the probability of fixation of neutral mutations. Mutations with selective effects shown in blue regions have an appreciable probability of fixation, while those shown in red are highly unlikely to fix (scale is displayed on right).

**Supp Figure 2:**
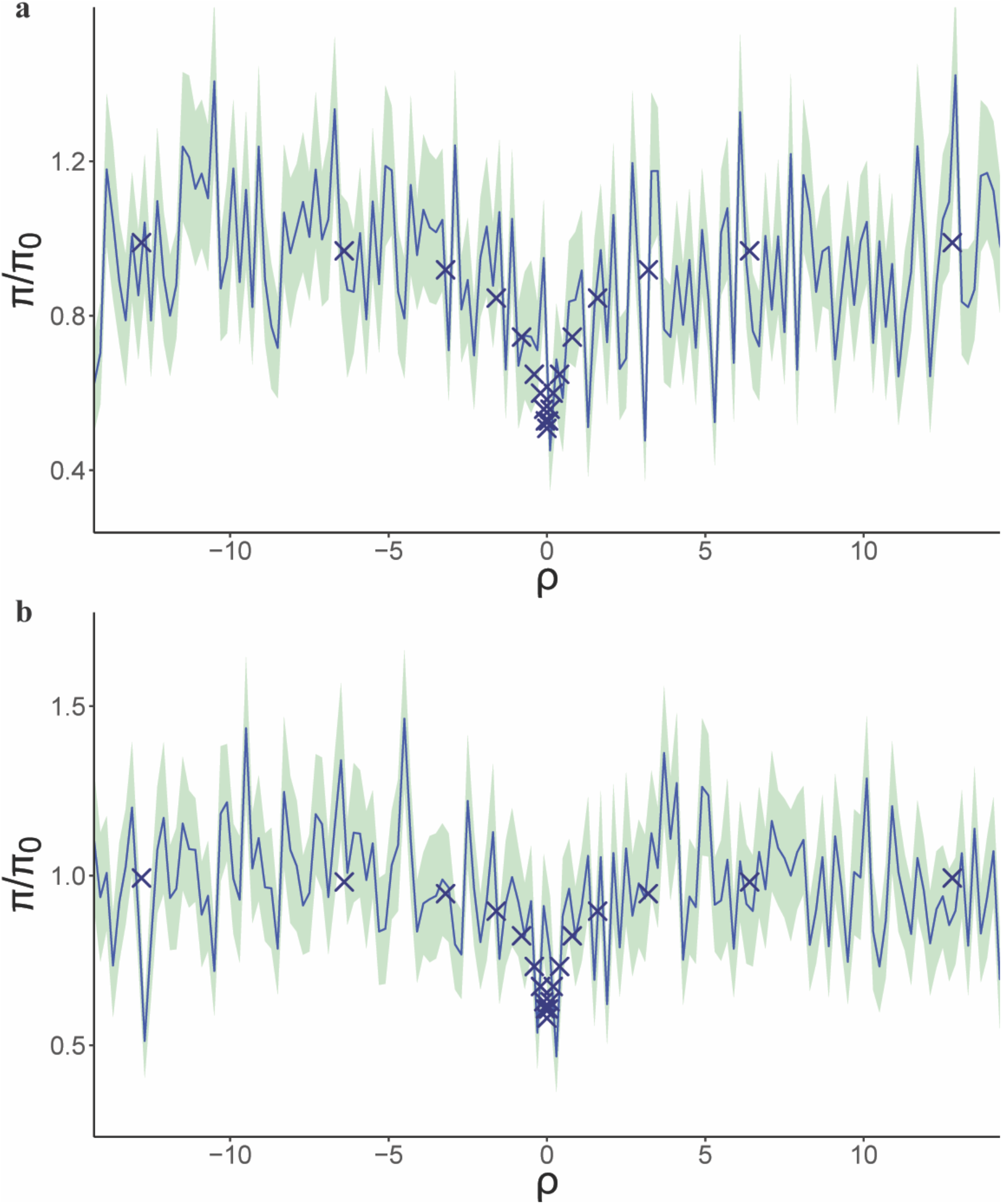
Neutral diversity relative to its mean value in the absence of selection, as a function of distance from the selected site, post-fixation of a mutation, with 2*Ns*_*a*_ = 5 and (a) *h* = 0 and (b) *h* = 1. Solid lines represent mean values of 100 replicates, shaded regions correspond to 1 SE above and 1 SE below the mean. Crosses correspond to simulations based on equations (27) of Tajima (1990).

**Supp Figure 3:**
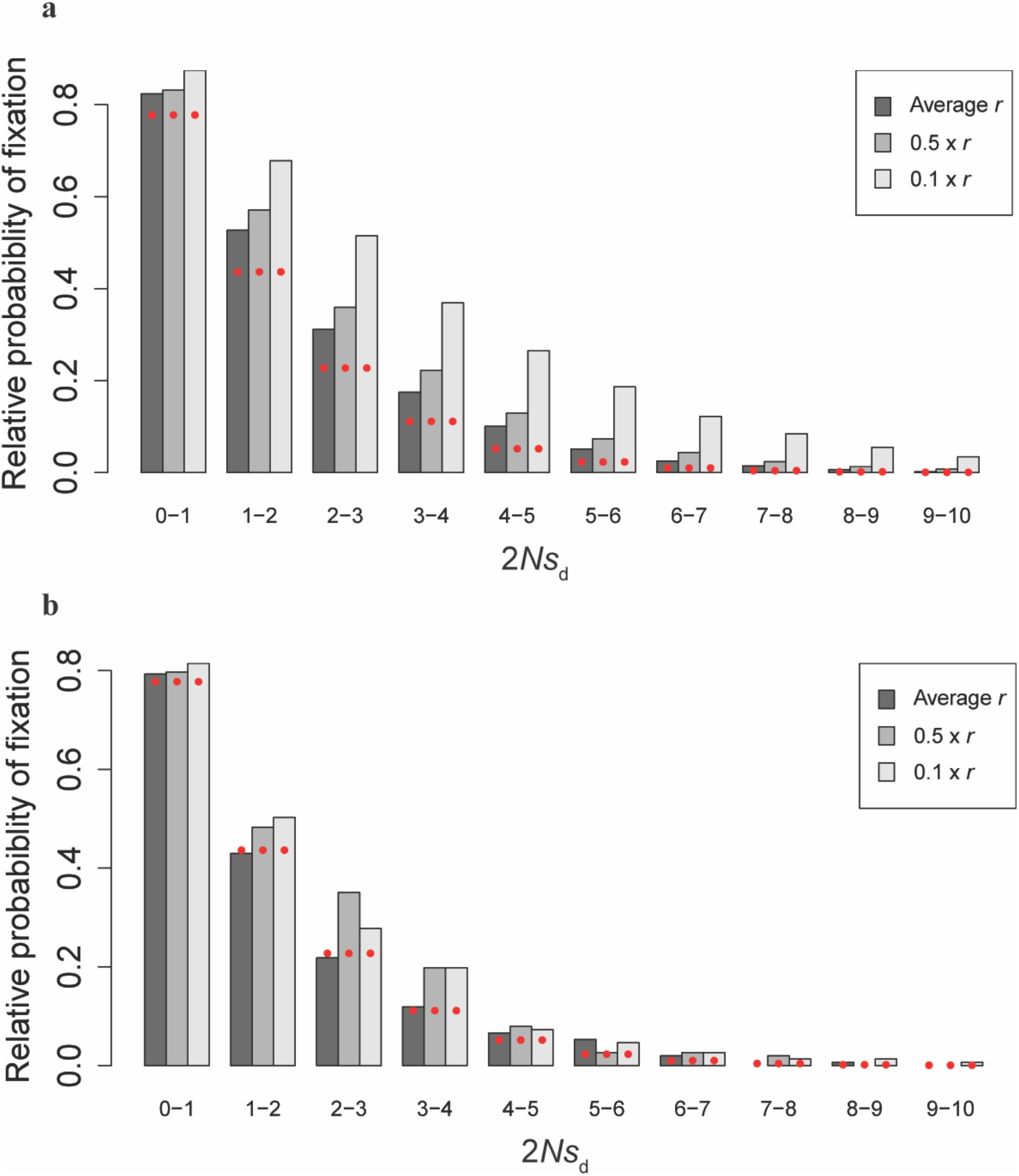
Frequencies of fixations of weakly deleterious mutations with possible interference amongst deleterious mutations relative to the probability of fixation of neutral mutations (*i.e*., 1/(2*N*)) for (a) *D. melanogaster* and (b) humans. Calculations of the probability of fixation from both simulations and theoretical expectations do not account for BGS. Red solid circles show the expected probabilities obtained using Equation 4.

**Supp Figure 4:**
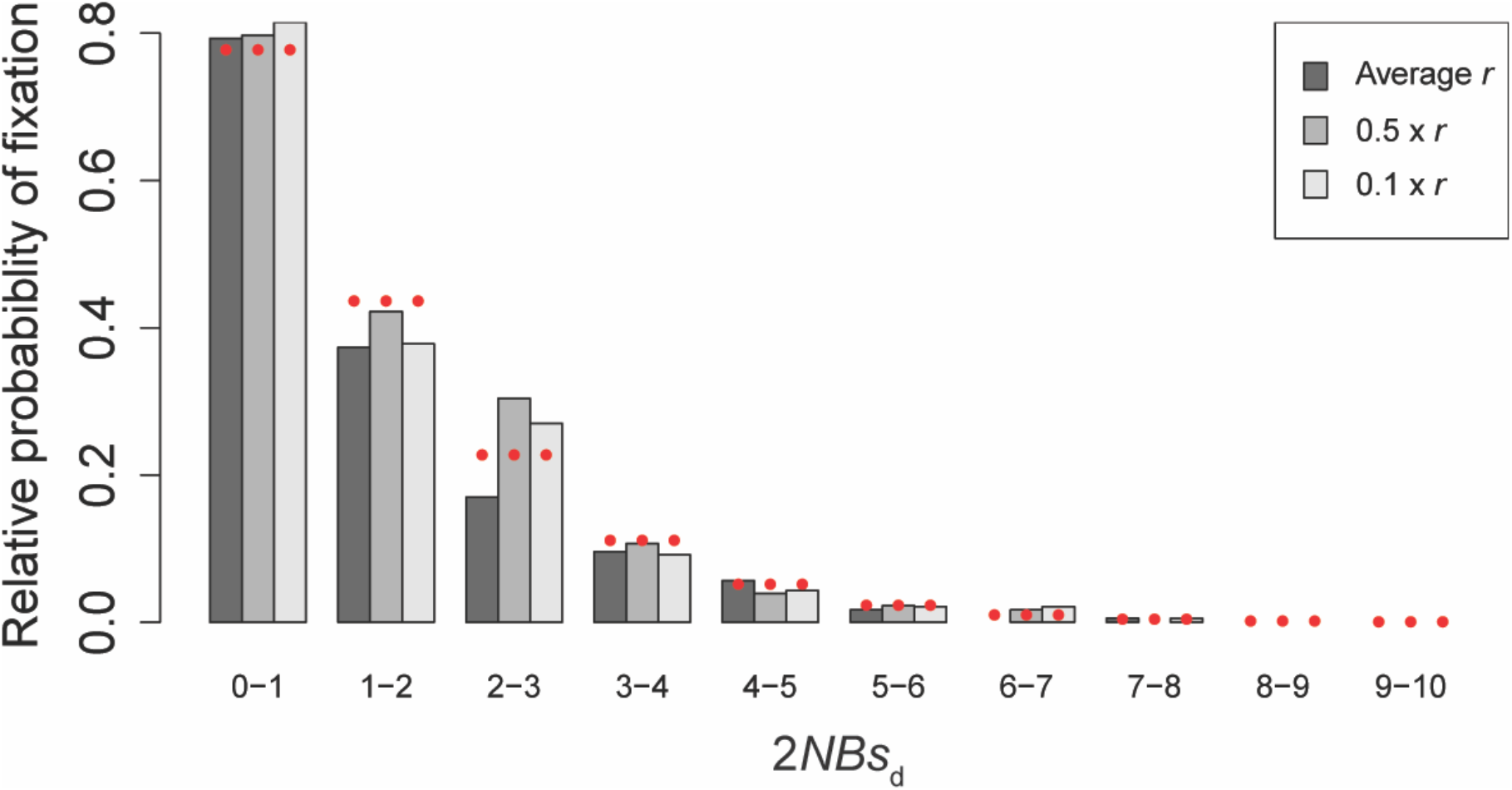
Frequencies of fixations of weakly deleterious mutations with possible interference amongst deleterious mutations relative to the probability of fixation of neutral mutations (*i.e*., 1/(2*N*)) for humans, when accounting for the effects of BGS. Red solid circles show the expected probability obtained by Equation 4.

**Supp Figure 5:**
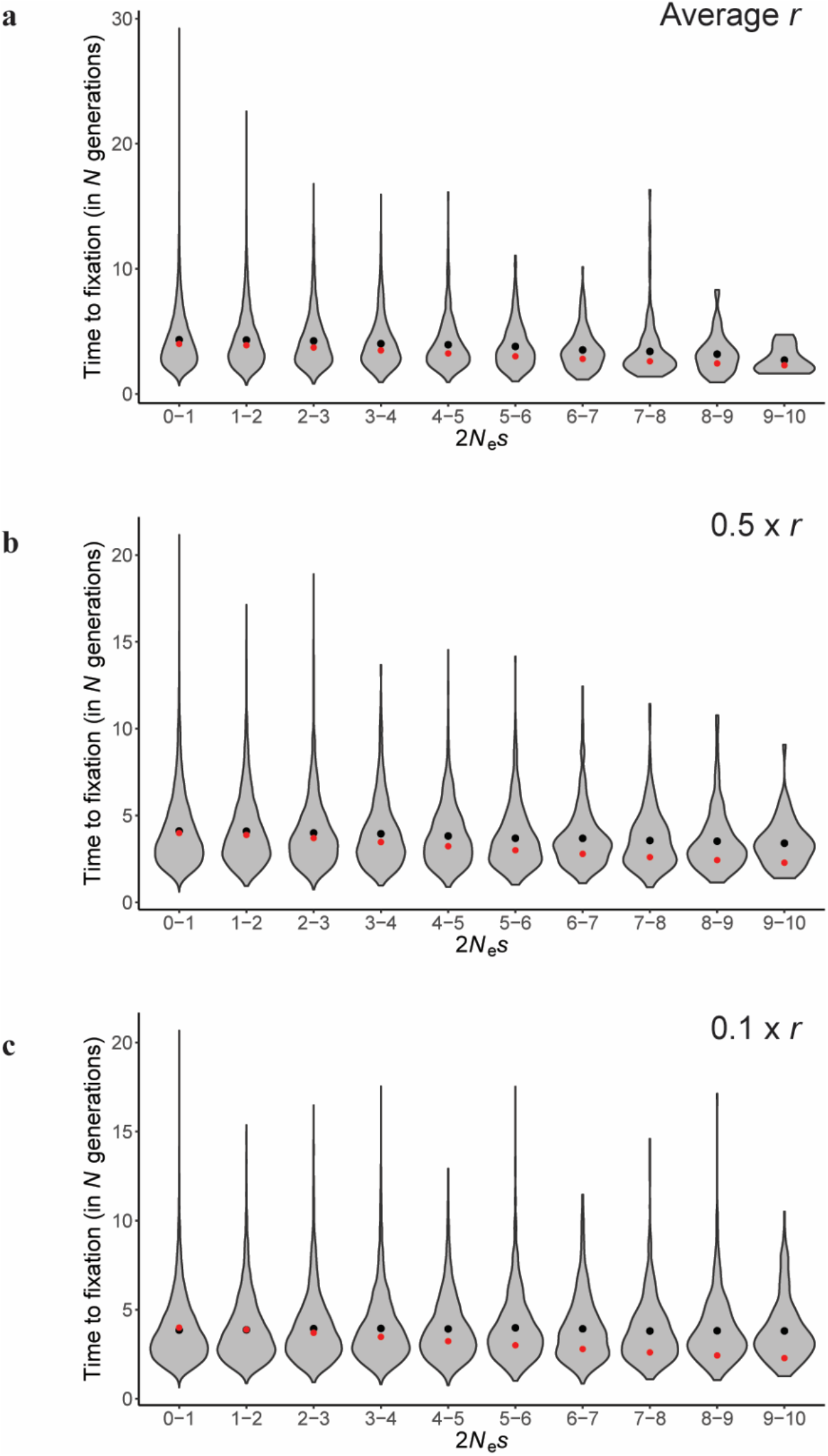
The distribution of the time to fixation (conditional on fixation) of mutations with possible interference amongst deleterious mutations in *D. melanogaster*. Red circles indicate the expected time to fixation (not accounting for interference or BGS) at the mid-point of the range of 2*N_e_s* shown, obtained by numerically integrating Equation 17 of Kimura & Ohta (1969).

**Supp Figure 6:**
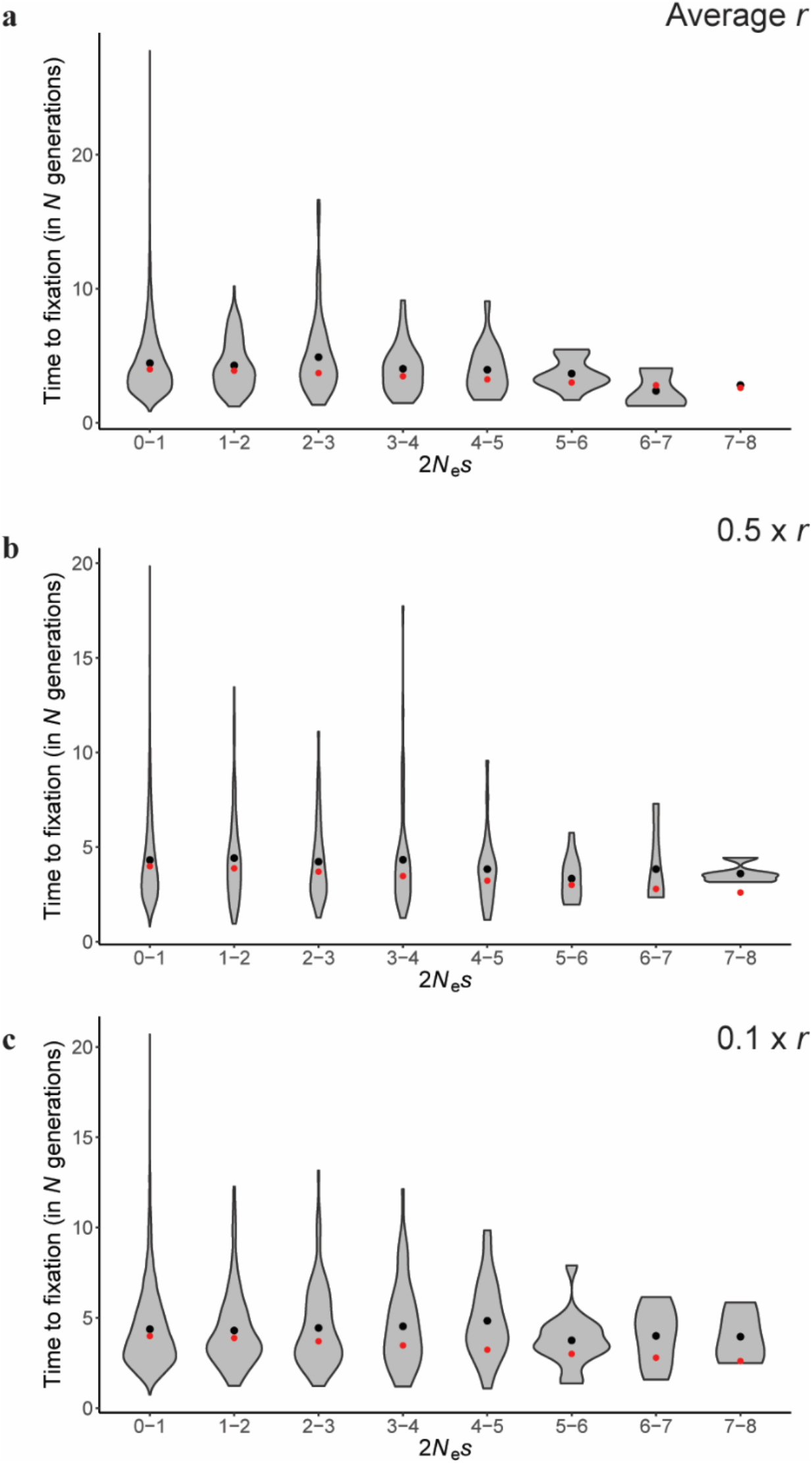
The distribution of the time to fixation (conditional on fixation) of mutations with possible interference amongst deleterious mutations in humans. Simulations were performed mimicking the exon-intron-intergenic structure of humans with the DFE at all exonic sites being: *f*_0_ = 0.51, *f*_1_ = 0.14, *f*_2_ = 0.14, and *f*_3_ = 0.21. Red circles indicate the expected time to fixation (not accounting for interference or BGS) at the mid-point of the range of 2*N_e_s* shown, obtained by numerically integrating Equation 17 of Kimura & Ohta (1969).

**Supp Figure 7:**
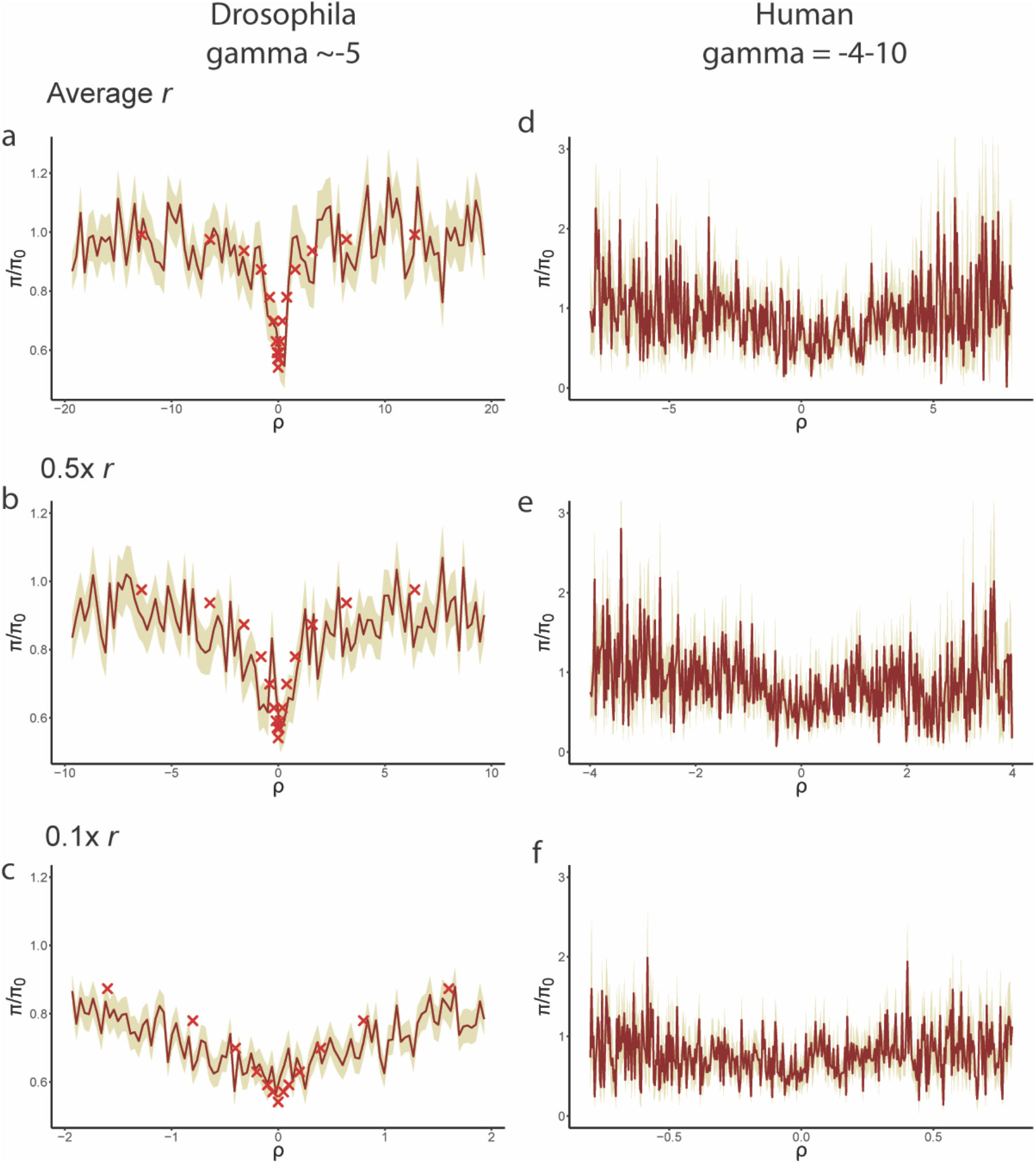
Neutral diversity relative to the mean value in the absence of selection, as a function of distance from the selected site, post-fixation of a mutation, with 2*Ns_d_* = 5 in the left panel corresponding to Drosophila-like parameters and 2*Ns_d_* between 4 and 10 in the right panel corresponding to human-like parameters. Solid lines represent mean values obtained from all observed substitutions, shaded regions correspond to 1 SE above and 1 SE below the mean. Crosses correspond to simulations based on equation (27) of Tajima (1990) for 2*Ns* = 5.

**Supp Figure 8:**
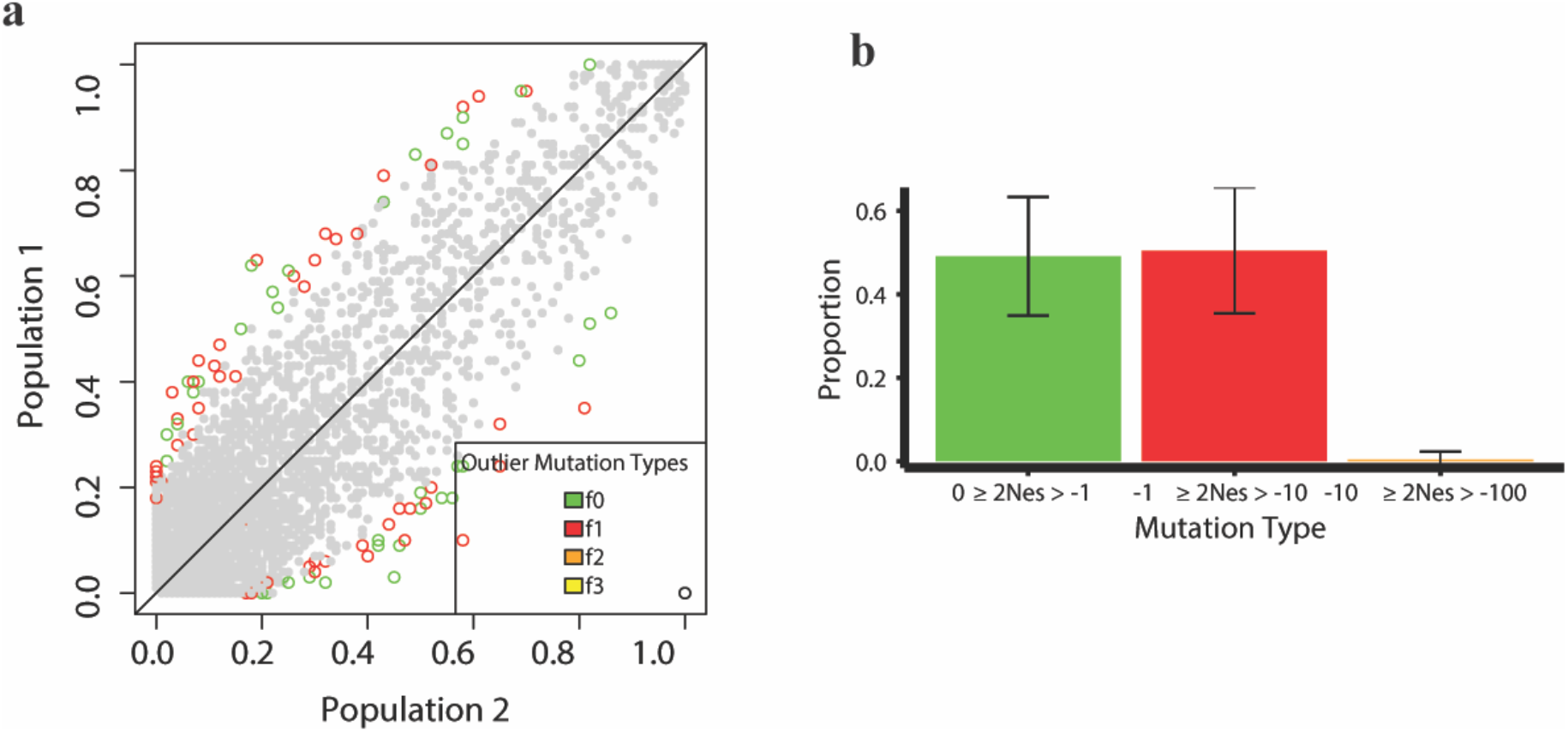
Allele frequencies of SNPs in European (population 1) *vs* African (population 2) populations - under the parameters inferred by Arguello *et al.* (2019), and where the selective effects of all mutations are not re-scaled after the split (thus selection is stronger in the African population). Simulated genomic elements experience purifying selection given by the DFE inferred in Johri *et al.* (2020). Colored open circles represent outliers. Green depicts effectively neutral mutations while warm colors depict deleterious mutations, with red representing weakly deleterious mutations*. Left panel*: Allele frequency plots are shown for 10 (out of 100) replicates. *Right panel*: The distribution of fitness effects of outlier mutations displaying the mean and standard deviation for all 100 replicates.

**Supp Figure 9:**
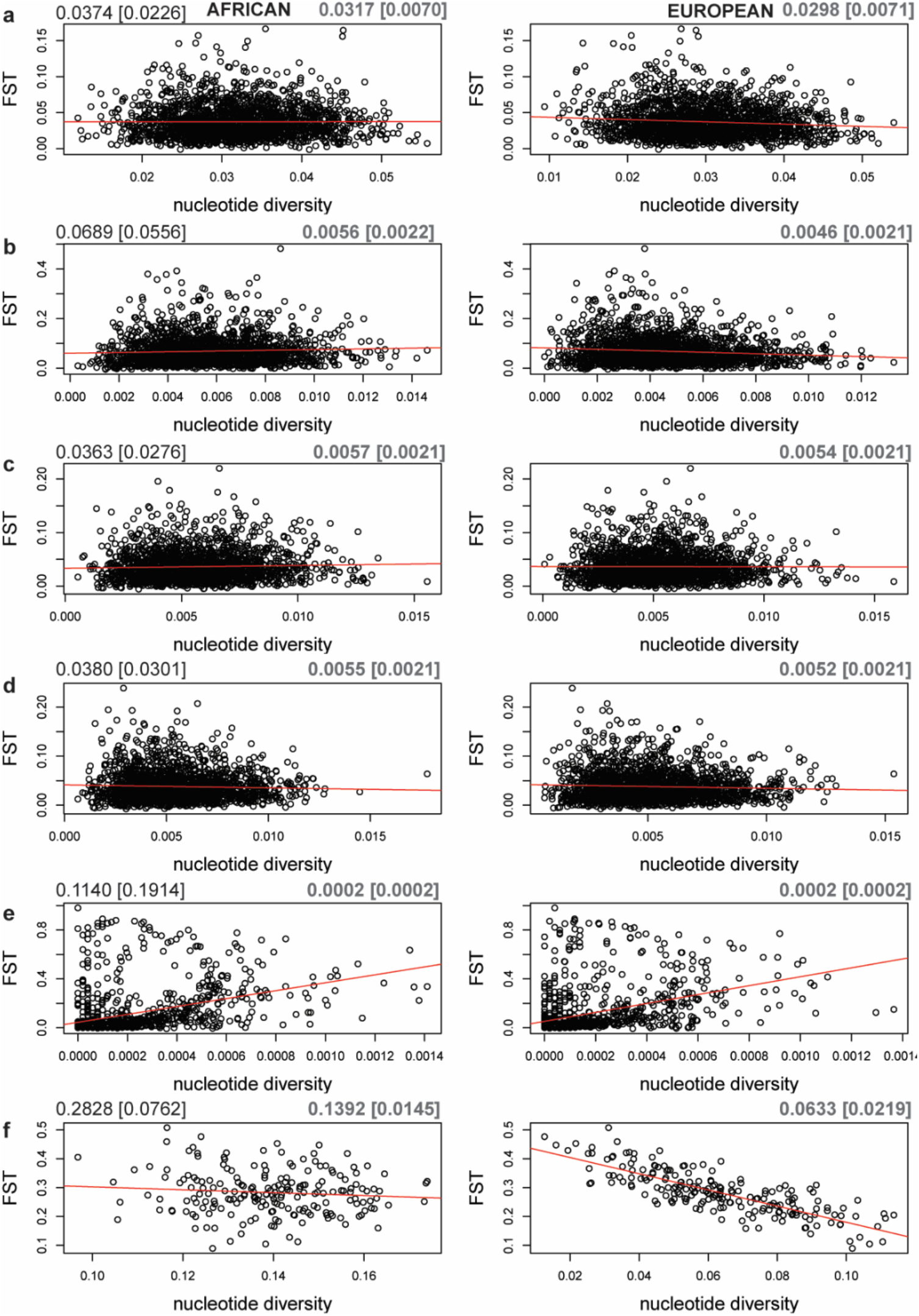
The relationship between *F_ST_* and nucleotide diversity calculated for neutral mutations alone, for the African and European populations simulated under six scenarios: (a) neutrality, (b) purifying selection (rescaled DFE), (c) purifying selection (not rescaled DFE), (d) purifying selection and weak positive selection, (e) purifying selection and strong positive selection, and (f) the neutral bottleneck of Li & Stephan 2006. In each panel, the upper left corner gives mean *F_ST_* values with its SD in parentheses. The upper right corners give the corresponding mean value of *π* with its SD in parentheses. The red line represents the best fitting linear regression. Note the difference in scale between scenarios.

**Supp Figure 10:**
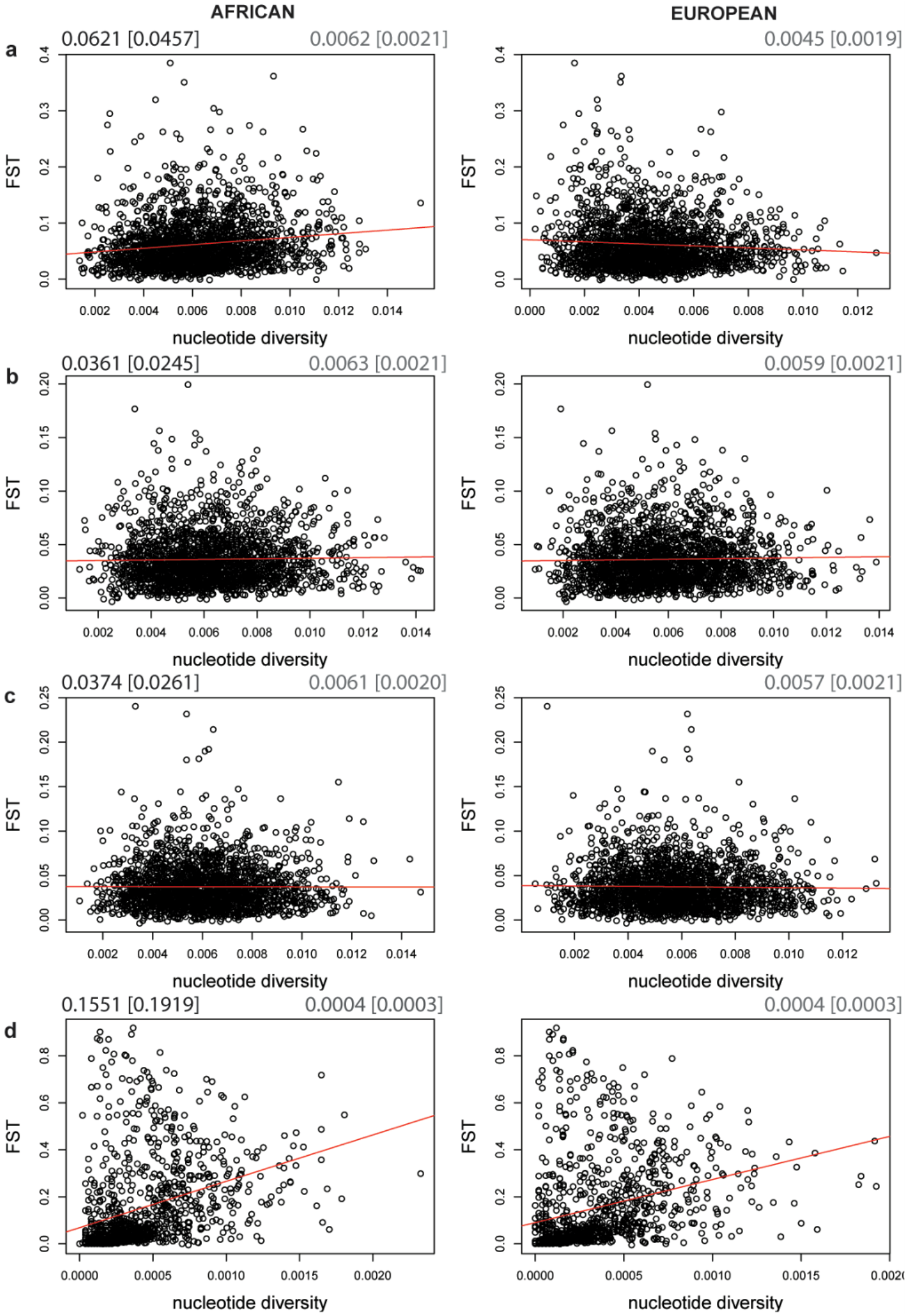
The relationship between *F_ST_* and nucleotide diversity calculated for weakly deleterious mutations alone, in the African and European populations simulated under four scenarios: (a) purifying selection (rescaled DFE), (b) purifying selection (not rescaled DFE), (c) purifying selection and weak positive selection, and (d) purifying selection and strong positive selection. On each, the upper left corner gives mean *F_ST_* values with its SD in parentheses. The upper right corner gives the corresponding mean value of *π* with its SD in parentheses. The red line represents the best fit linear regression. Note the difference in scale between scenarios.

**Supp Figure 11:**
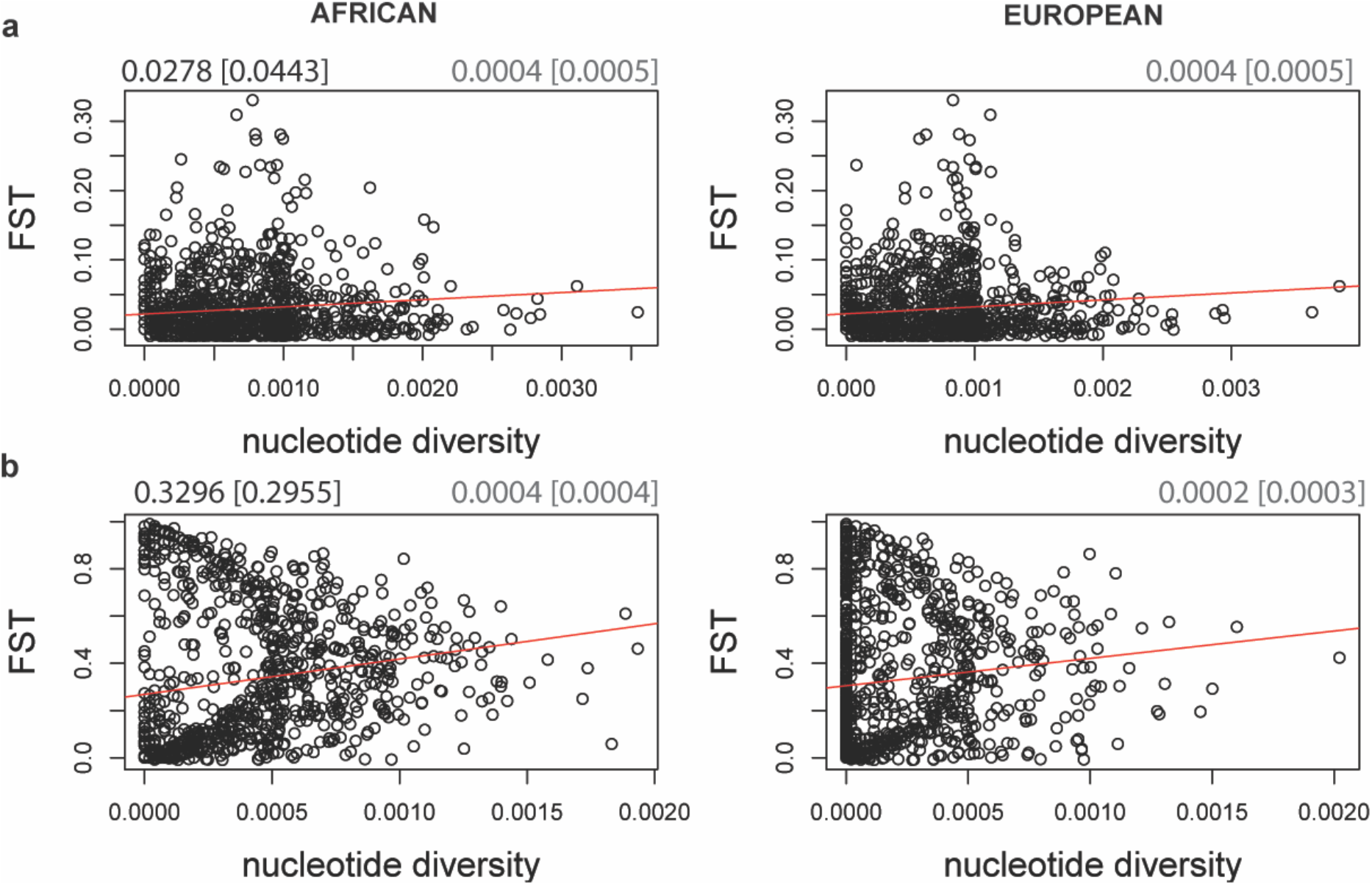
The relationship between *F_ST_* and nucleotide diversity calculated for only beneficial mutations in the African and European populations simulated under two scenarios: (a) purifying selection and weak positive selection, and (b) purifying selection and strong positive selection. On each, the upper left corner gives mean *F_ST_* values with its SD in parentheses. The upper right corner gives the corresponding mean value of *π* with its SD in parentheses. The red line represents the best fit linear regression. Note the difference in scale between scenarios.

